# Bacterial Hsp90 promotes virulence factor production through maintenance of NRPS megaenzymes

**DOI:** 10.1101/2025.07.01.662645

**Authors:** Alison Fratacci, Amine Ali Chaouche, Marie Corteggiani, Flora Honoré, Manel Khelil Berbar, Nadège Bossuet-Greif, Sébastien Dementin, Sophie Bleves, Maya Belghazi, Régine Lebrun, Jean-Philippe Nougayrède, Eric Oswald, Bérengère Ize, Olivier Genest

**Author notes:** Co-corresponding authors: Olivier Genest and Bérengère Ize: Olivier Genest, CNRS, BIP UMR 7281, 31 Chemin Joseph Aiguier, 13402 Marseille, Cedex 20, France. Bérengère Ize, CNRS, LISM UMR 7255, IMM, 31 Chemin Joseph Aiguier, 13402 Marseille, France. These authors contributed equally to this work (Alison Fratacci, Amine Ali Chaouche). **Author Contributions:** Designed research: AF, AAC, RL, SB, JPN, EO, BI, OG; Performed research: AF, AAC, MC, FAH, MKB, NBG, MB, RL, JPN, BI, OG; Analyzed data: AF, AAC, MC, FAH, MKB, NBG, SD, SB, MB, RL, JPN, EO, BI, OG; Wrote the paper: BI and OG with contributions from all the authors; Funding acquisition: RL, SB, JPN, EO, BI, OG; Supervised the study: BI, OG.

## Abstract

Pathogenic bacteria produce virulence factors critical to host infection. Here, we demonstrate the crucial role of the bacterial Hsp90 chaperone in the production of two major virulence factors, the colibactin genotoxin in *Escherichia coli* and the pyoverdine siderophore in *Pseudomonas aeruginosa*. Colibactin, a hybrid polyketide/non-ribosomal peptide (PK-NRP), and pyoverdine, a non-ribosomal peptide (NRP), are metabolites produced by complex biosynthetic pathways involving large cytoplasmic enzymes called megasynthases. Using comparative proteomics, we found that megasynthase abundance was markedly reduced in *hsp90* deletion mutants of *E. coli* and *P. aeruginosa* compared to wild-type strains. This reduction was independent of transcriptional or translational regulation. We further revealed an interplay between Hsp90 and the HslUV protease in controlling megasynthase levels. Remarkably, we found that Hsp90 stabilizes additional NRP and PK-NRP megasynthases, suggesting a general role for Hsp90 as a chaperone of these enzymes. These findings open new avenues for enhancing the biosynthesis of complex metabolites for biotechnological applications through proteostasis modulation, and may also have implications for combating bacterial infections.

## Introduction

Bacteria possess an arsenal of adaptive pathways to cope with the adverse conditions encountered in the constantly changing and stressful environment in which they evolve (1–6). One example illustrating this is bacterial infection, during which bacteria face host defenses such as increased temperature, oxidative and nitrosative stresses, as well as depletion of some essential elements like iron (7, 8). Among the many bacterial responses that are implemented, a complex network of chaperones and proteases maintains protein homeostasis to circumvent the deleterious effects of stress on protein folding, stability, and activity (9–14).

Hsp90 is an ATP-dependent chaperone found in bacteria and eukaryotes (15–18). It is a V-shaped dimer made of three domains, a N-terminal domain that hydrolyzes ATP together with the middle domain, and a C-terminal domain responsible for dimerization (19–23). Its chaperone cycle depends on ATP binding and hydrolysis, leading to a sequence of large conformational changes conserved in bacteria and eukaryotes (24). Many cochaperones modulate these conformational changes in eukaryotes, whereas no Hsp90 cochaperone has been identified to date in prokaryotes (16, 25, 26). Hundreds of clients of eukaryotic Hsp90 have been discovered including oncoproteins, leading to the development of a large number of Hsp90 specific inhibitors (27–29).

Although only a few clients of bacterial Hsp90 (also known as HtpG) have been identified, it is essential under stress conditions in some bacteria (*Synechococcus sp. PCC 7942*, *Shewanella oneidensis*) (30–33), whereas its presence seems to be dispensable for growth in others such as *E. coli* (34). Importantly, bacterial Hsp90 is associated with the pathogenesis and virulence of pathogens including *Salmonella Typhimurium* (35), *Leptospira interrogans* (36), *Edwardsiella tarda* (37), and extra intestinal pathogenic *E. coli* (ExPEC) (38). While the exact role of Hsp90 in most of these organisms is still unknown, Garcie *et al* showed that it is involved in the biosynthesis of the colibactin toxin and the yersiniabactin siderophore in ExPEC, although the molecular mechanism for this effect is still lacking (38). Colibactin and yersiniabactin are secondary metabolites that belong to the family of hybrid polyketide non-ribosomal peptides (PK-NRP) (39–41). Interestingly, it has been reported that Hsp90 participates in the production of other hybrid PK-NRP or NRP compounds like the antibiotic albicidin in *Xanthomonas albilineans* (42), the biosurfactant arthrofactin in *Pseudomonas sp. MIS38* (43), and the siderophore pyoverdine in *P. aeruginosa* (44). The production of these molecules depends on complex pathways often involving dozens of proteins which allow the cytoplasmic biosynthesis of a precursor, its export across the inner membrane, its periplasmic maturation and its secretion into the external environment. Given the role of these natural molecules in bacterial virulence and their importance for biotechnological applications, it is of great interest to discover how Hsp90 participates in these biosynthetic pathways.

Here, we demonstrate the crucial role of Hsp90 in the biosynthesis of hybrid PK-NRP and NRP metabolites. Focusing on the production of two major bacterial virulence factors, colibactin in *E. coli* and pyoverdine in *P. aeruginosa*, we find that the amount of large enzymes called megasynthases, involved in the cytoplasmic biosynthesis of these virulence factors is dramatically decreased in the absence of Hsp90. Hsp90 acts at the posttranslational level, and an interplay occurs between Hsp90 and the HslUV protease to regulate the levels of these megasynthases. We further show that Hsp90 maintains the level of megasynthases required for the biosynthesis of other NRP or PK-NRP molecules, suggesting a conserved role for Hsp90 in the production of these molecules is in bacteria.

## Results

### Hsp90 stabilizes large enzymes involved in colibactin production in *E. coli*

Colibactin is a genotoxin produced by Extra intestinal pathogenic *E. coli* (ExPEC) and commensal strains, including the *E. coli* M1/5 strain (45). Colibactin belongs to the family of hybrid polyketide (PK) non-ribosomal peptide (NRP) and is produced by a complex biosynthetic pathway involving 19 proteins encoded in the *pks* island (41, 46–48) (**Fig. 1A**). A colibactin precursor is synthetized in the cytoplasm by an assembly line made of large enzymes, called megasynthases (ClbN, B, C, H, I, J, K and O, from 89 kDa to 352 kDa) that either belong to the family of non-ribosomal peptide synthetases (NRPS), polyketide synthases (PKS), or hybrid NRPS and PKS (NRPS-PKS), as well as additional ‘accessory’ proteins (ClbA, D, E, F, G, Q, L, and S). The precursor is then translocated to the periplasm through ClbM, activated by the peptidase ClbP, and secreted. An immunity protein, ClbS, is found in the cytoplasm to prevent self-DNA damage. Interestingly, Hsp90_Ec_ (Hsp90 of *E. coli*) was shown to be required for colibactin synthesis (38), yet, its specific function remains unknown.

**Fig. 1.**
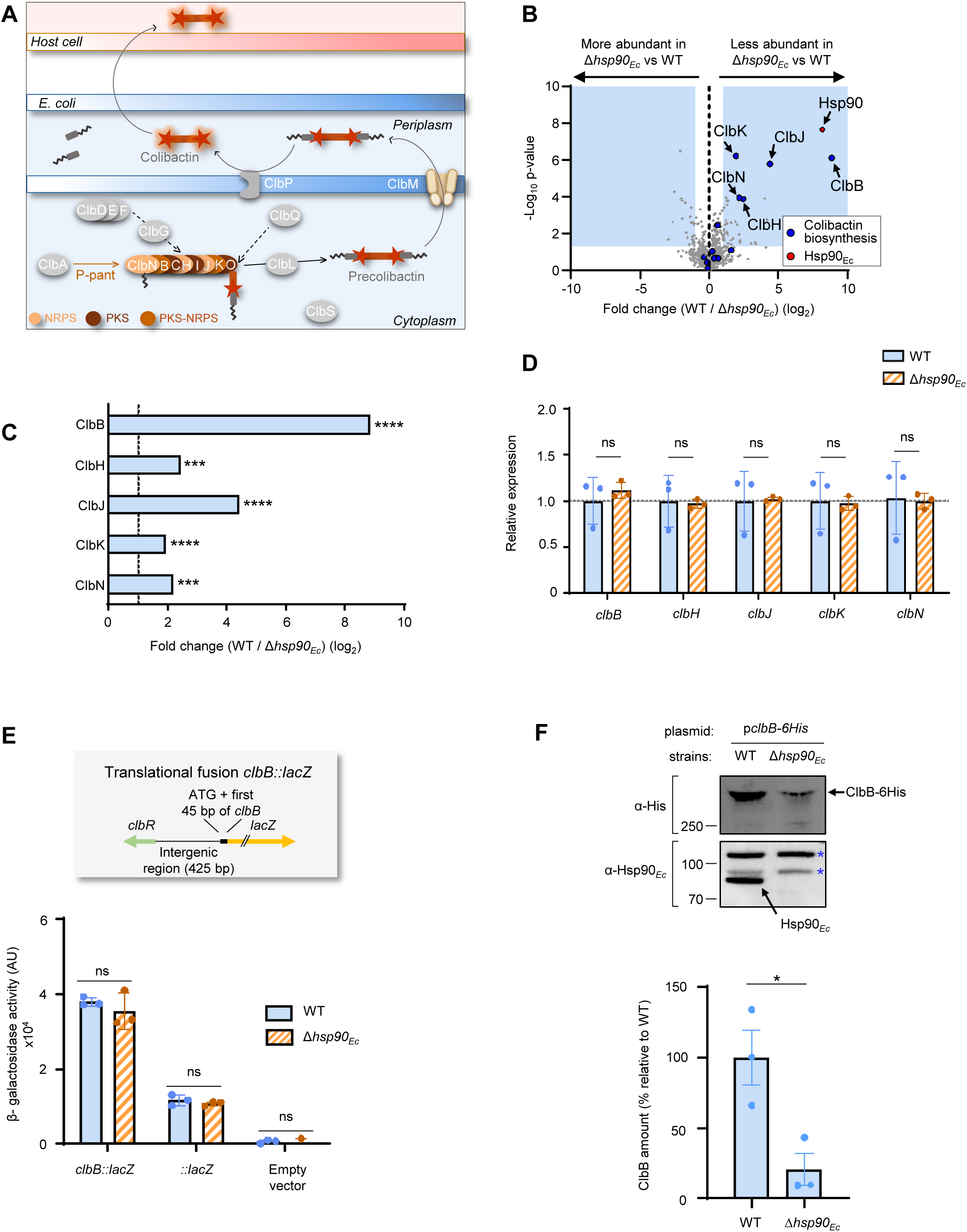
Hsp90_Ec_ acts at the post-translational level to maintain the abundance of megasynthases involved in colibactin production in *E. coli*. (A). Schematic representation of the biosynthesis pathway of colibactin. Colibactin is a hybrid non-ribosomal peptide-polyketide toxin. An inactive form, pre-colibactin is synthesized in the cytoplasm by an assembly line composed of large enzymes called NRPS (ClbH, ClbJ, ClbN), PKS (ClbC, ClbI, and ClbO), or hybrid NRPS-PKS (ClbB, ClbK), as well as other proteins (gray). After being transported in the periplasm, pre-colibactin is activated by a peptidase, and can reach the host cell causing DNA damages. (B) Global comparative quantitative proteomics of *E. coli* WT and Δ*hsp90* strains. The protein content of the two strains grown at 37°C was analyzed by LFQ mass spectrometry, and the relative abundance of proteins is reported. The X axis indicates the protein fold change (log_2_) between the two strains and the Y axis indicates the p-value (-log_10_) of this difference. Regions of the graph framed in light blue indicate proteins whose abundance varied significantly between the two strains (fold change >2; p-value <0.05). Proteins of interest are pointed by arrows. Hsp90_Ec_ is colored in red, and proteins involved in colibactin biosynthesis in blue. The experiments were performed with 4 biological replicates for each strain. (C) Relative abundance of megasynthases involved in colibactin biosynthesis measured in panel B. p-values associated with each ratio indicate that the differences are significant (****P ≤ 0.0001, ***P ≤ 0.001). (D) Hsp90*_Ec_* does not modify expression of the genes encoding the megasynthases. mRNA from *E. coli* WT and Δ*hsp90_Ec_* strains was extracted, retro-transcribed in cDNA, and quantitative-PCR was performed using specific oligonucleotides to amplify *clbB*, *clbH*, *clbJ*, *clbK*, and *clbN*. Data are shown as fold variation of expression in the Δ*hsp9*0*_Ec_* strain relative to the WT strain. Dot line (y=1) indicates the relative expression value for which there is no variation compared to the expression in the WT strain. Data from three biological replicates are shown as mean ± SEM. (E) Hsp90*_Ec_* does not modify translation of the ClbB protein. A 686 bp sequence including *clbR*, the promoter of *clbB*, RBS, ATG, and the first 45 nucleotides of *clbB* was cloned upstream of the *lacZ* gene (from amino acid 9) in a pACYC184 vector. The *clbR* gene encodes a transcriptional regulator located upstream of *clbB*. The resulting plasmid was introduced in the WT and Δ*hsp90_Ec_* strains. Plasmid with *lacZ* only (without promoter) and an empty vector were used as controls. β-galactosidase activity that reflects the level of translation of ClbB was measured. Data from three biological replicates are shown as mean ± SEM. (F) Top panel: ClbB with a 6-His tag (353.4 kDa) was produced from a plasmid under the control of an arabinose promoter in WT and Δ*hsp9*0*_Ec_* strains. ClbB was detected by immunoblot with a anti 6His antibody and Hsp90*_Ec_* with an anti-Hsp90*_Ec_* antibody. Note that two additional bands indicated by stars are non-specifically detected by the Hsp90*_Ec_* antibody and are used as loading control to show that the same amount of proteins was loaded in each well. The Western blots shown are representative of three independent experiments. Molecular weight markers (kDa) are indicated on the left. Bottom panel: the intensity of the bands corresponding to ClbB was quantified using ImageJ. Data from three biological replicates are shown as mean ± SEM. In D, E and F, results from t-tests indicate whether the differences measured are significant (*: p-value ≤ 0.05) or not (ns).

Since some Hsp90 clients have been proposed or shown to be degraded in the absence of Hsp90 (33, 38, 49, 50), we conducted a global comparative and quantitative proteomic analysis between *E. coli* wild-type (WT) and a Δ*hsp90* mutant strain, deleted of the *htpG* gene that encodes Hsp90. We hypothesized that the mutant strain would exhibit reduced levels of Hsp90 clients. Total proteomes of the two strains grown under hypoxic conditions - known to favor colibactin production - were analyzed using label-free quantitation (LFQ) mass spectrometry (**Fig. 1B**). In comparison to the WT strain, the levels of 25 more abundant proteins and 38 less abundant proteins were found to be statistically significant in the absence of Hsp90_Ec_ (fold change>2 (Log2>1), p-value<0.05) (**Datasets S1 and S2**). Interestingly, among the 38 proteins whose amount was significantly reduced in the absence of Hsp90_Ec_, 5 proteins (ClbB, ClbH, ClbJ, ClbK, and ClbN) involved in colibactin biosynthesis were identified (**Fig. 1B and 1C**). These five proteins are part of the cytoplasmic assembly line of colibactin biosynthesis, and are classified either as NRPS (ClbH, ClbJ, ClbN), or hybrid NRPS-PKS (ClbB, ClbK) (**Fig. 1A**). Among the 19 proteins encoded in the *pks* island and involved in colibactin biosynthesis, 10 additional proteins were detected in the proteomic analysis, however their abundance did not vary much between the WT and Δ*hsp90*_Ec_ strains (**Fig. 1B, S1A**).

To rule out the possibility that the reduced abundance of ClbB, ClbH, ClbJ, ClbK, and ClbN is due to an indirect effect of *hsp90*_Ec_ deletion on the transcription of the corresponding *clb* genes, qRT-PCRs were performed on WT and Δ*hsp90*_Ec_ strains grown under the conditions of the proteomic experiments. Notably, we found that expression of *clbB*, *clbH*, *clbJ*, *clbK* and *clbN* did not significantly vary between the two strains (**Fig. 1D**). Next, to test for a putative role of Hsp90_Ec_ in the translational regulation of the Clb proteins, a plasmidic *clbB*::*lacZ* translational fusion was introduced in WT and Δ*hsp90 E. coli* strains. We chose ClbB because it was the protein with the highest fold reduction in the absence of Hsp90_Ec_. No significant difference in β-galactosidase activity was measured between the two strains, indicating that the absence of Hsp90_Ec_ did not modify transcription and translation of *clbB* (**Fig. 1E**).

Finally, to assess whether Hsp90 regulates the Clb proteins at a post-translational level, we placed *clbB* with a sequence encoding a hexa-histidine (6His) epitope-tag under the control of an arabinose-inducible promoter to remove the physiological transcriptional and translational regulation of *clbB*. The functionality of ClbB-6His was evaluated, and no alteration was observed (**Fig. S1B**). Notably, upon induction, ClbB-6His was present in a significantly lower amount in the Δ*hsp90*_Ec_ mutant in comparison to the WT cells or to the complemented Δ*hsp90*_Ec_ mutant carrying a plasmid allowing Hsp90_Ec_ production (**Fig. 1F and S1C**).

Altogether, these experiments demonstrate that the reduction of the colibactin toxin observed in the absence of Hsp90_Ec_ (38) results from the decrease at the post-translational level of the amount of several NRPS and NRPS-PKS proteins involved in colibactin biosynthesis.

### Hsp90 participates in the biosynthesis of the NRP pyoverdine in *P. aeruginosa*

In addition to colibactin, it has been suggested that Hsp90 plays a role in the production of another NRP molecule, the siderophore pyoverdine secreted by the opportunistic pathogen *P. aeruginosa* (44). Pyoverdine, a major virulence factor of *P. aeruginosa*, is a non-ribosomal peptide molecule produced under iron-limiting conditions by a complex biosynthetic pathway involving about thirty proteins (51, 52). A pyoverdine precursor is synthesized by a putative cytoplasmic complex called siderosome constituted by 4 NRPS (PvdD, PvdI, PvdJ and PvdL) with molecular weights ranging from 240 kDa to 569 kDa, as well as additional proteins (PvdH, PvdF, PvdA) (**Fig. 2A**) (53–55). The pyoverdine precursor is then translocated to the periplasm where it is modified by several additional proteins to form the fluorescent pyoverdine before being secreted into the external environment. There, pyoverdine chelates iron, and the iron-bound siderophore can be internalized by *P. aeruginosa* to increase intracellular iron levels.

**Fig. 2.**
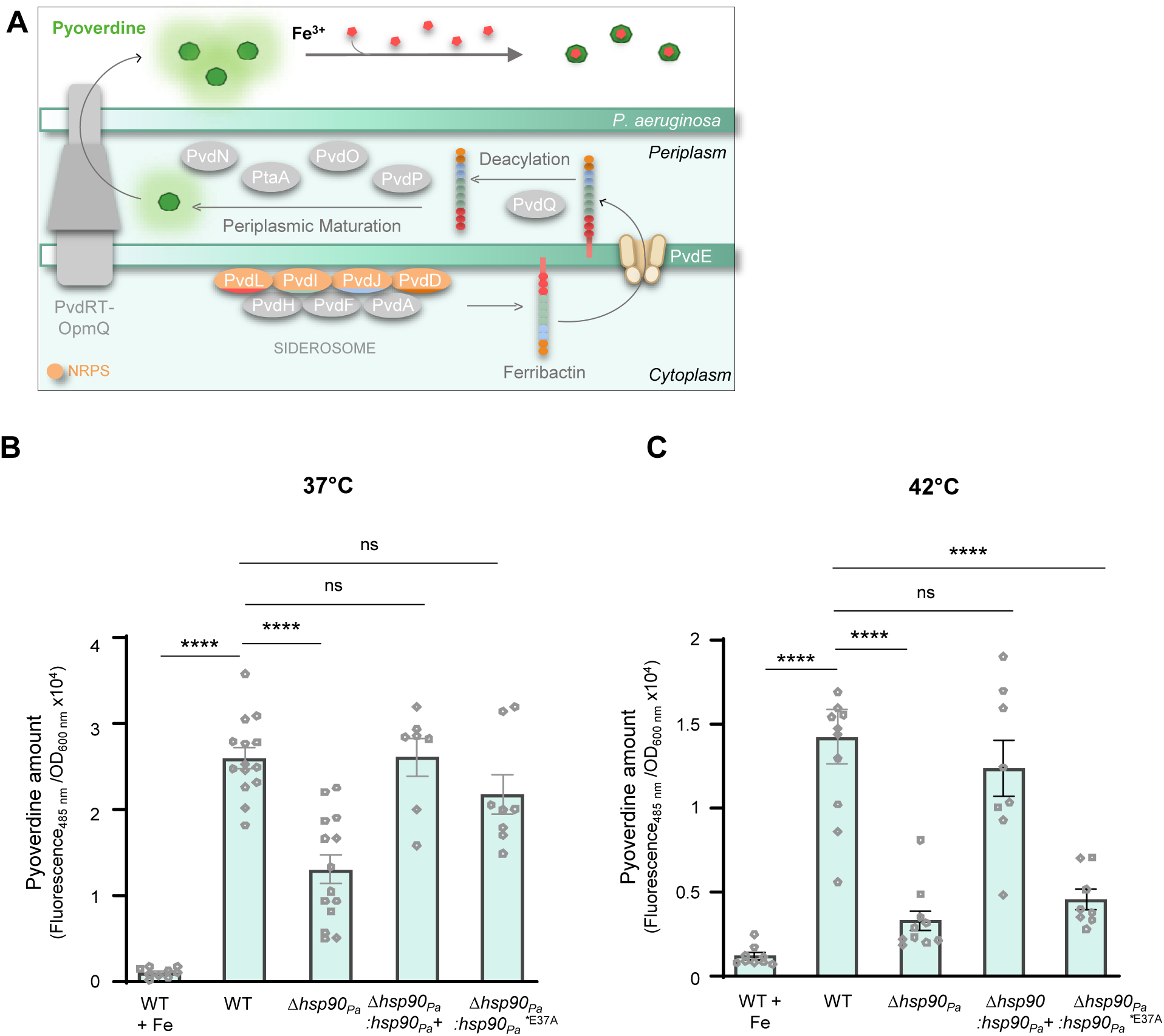
Hsp90*_Pa_* participates in pyoverdine production in *P. aeruginosa*. (A) Schematic representation of the biosynthetic pathway of the siderophore pyoverdine. The pyoverdine precursor, ferribactin is synthesized in the cytoplasm by an assembly line made of four NRPS (PvdL, PvdI, PvdJ, and PvdD) and three associated proteins (PvdH, PvdF, and PvdA). These proteins form a complex called the siderosome. Ferribactin is subsequently translocated to the periplasm where it is activated by several enzymes before being secreted in the external medium where it interacts with Fe^3+^. Iron can then be imported back into the cell by additional proteins not shown on this schematic representation. (**B** and **C**) Pyoverdine production measurement. *P. aeruginosa* WT, Δ*hsp90*, Δ*hsp90*:*hsp90*+, and Δ*hsp90*:*hsp90*\*_E37A_ strains were grown at 37°C (B) or 42°C (C) in iron-deficient medium (CAA). As a control, iron was supplemented (WT+Fe). After 5h growth, pyoverdine was quantified by measuring fluorescence (excitation: 400 nm; emission: 485 nm) that was standardized to the number of cells (OD_600_). Data from at least seven biological replicates are shown as mean ± SEM. Results of one-way ANOVA tests indicate whether the differences measured are significant (****: p-value ≤ 0.0001; ***: p-value ≤ 0.001) or not significant (ns, p-value > 0.05).

To robustly confirm the effect of Hsp90 on pyoverdine production, WT, Δ*hsp90*, and a Δ*hsp90*:*hsp90*+ complemented strain of *P. aeruginosa* were grown in iron-limiting conditions (CAA) at various temperatures, and pyoverdine production was measured after 5 hours of growth from culture supernatants using the natural fluorescence of this molecule. Note that the absence of Hsp90 from *P. aeruginosa* (Hsp90_Pa_) did not alter bacterial growth in the tested conditions (**Fig. S2A**). Although similar levels of fluorescence were observed at 30°C between WT and Δ*hsp90*_Pa_ strains (**Fig. S2B**), a 1.6-time reduction in pyoverdine fluorescence was observed at 37°C in the Δ*hsp90*_Pa_ strain compared to WT (**Fig. 2B**). Interestingly, when strains were grown under heat stress (42°C), a 6-time decrease of pyoverdine fluorescence was measured in the absence of Hsp90_Pa_ compared to WT, and this defect was restored in the complemented strain (Δ*hsp90*_Pa_::*hsp90* _Pa_ +) (**Fig. 2C**).

To determine whether the ATP-dependent chaperone activity of Hsp90_Pa_ is important for pyoverdine production, the glutamate residue at position 37, homologous to the catalytic glutamate 34 in Hsp90_Ec_, was substituted to alanine to abolish ATPase activity (56, 57). When produced in the Δ*hsp90 P. aeruginosa* strain, Hsp90_Pa_E37A did not support the production of pyoverdine, even though production of the variant protein was increased compared to WT (**Fig. 2B, 2C, and S2C**). These results show that pyoverdine production in *P. aeruginosa* requires the ATP-dependent activity of Hsp90, and that the importance of Hsp90 increases with temperature as expected for a chaperone whose role is usually more important under heat stress.

### Hsp90 stabilizes NRP enzymes involved in pyoverdine biosynthesis in *Pseudomonas aeruginosa*

To identify potential clients of Hsp90_Pa_ involved in pyoverdine production, the proteomes of *P. aeruginosa* WT and Δ*hsp90* strains grown under iron-limiting conditions at 37°C and 42°C were compared by LFQ mass spectrometry. Interestingly, these experiments revealed that the 4 NRPS proteins involved in the biosynthesis of pyoverdine (**Fig. 2A**) were less abundant in the absence of Hsp90_Pa_. Indeed, PvdD, PvdI, PvdJ and PvdL were decreased between 1.9- and 3.2-fold at 37°C (**Fig. 3A and C**) and the reduction in abundance was even more pronounced at 42°C, ranging between 2.3 and 15.4-fold (**Fig. 3B and C**). PvdJ exhibited the most significant reduction in abundance in the absence of Hsp90_Pa_ (**Fig. 3A-C**). It is important to note that the abundance of other proteins involved in pyoverdine biosynthesis (non NRPS) and detected in the proteomic experiments, was not significantly reduced in the Δ*hsp90*_Pa_ strain at 37°C and 42°C (**Fig 3A, 3B, and S3A**). Additional proteins whose abundance significantly varied between the two strains are listed in **Datasets S3-S6**.

**Fig. 3.**
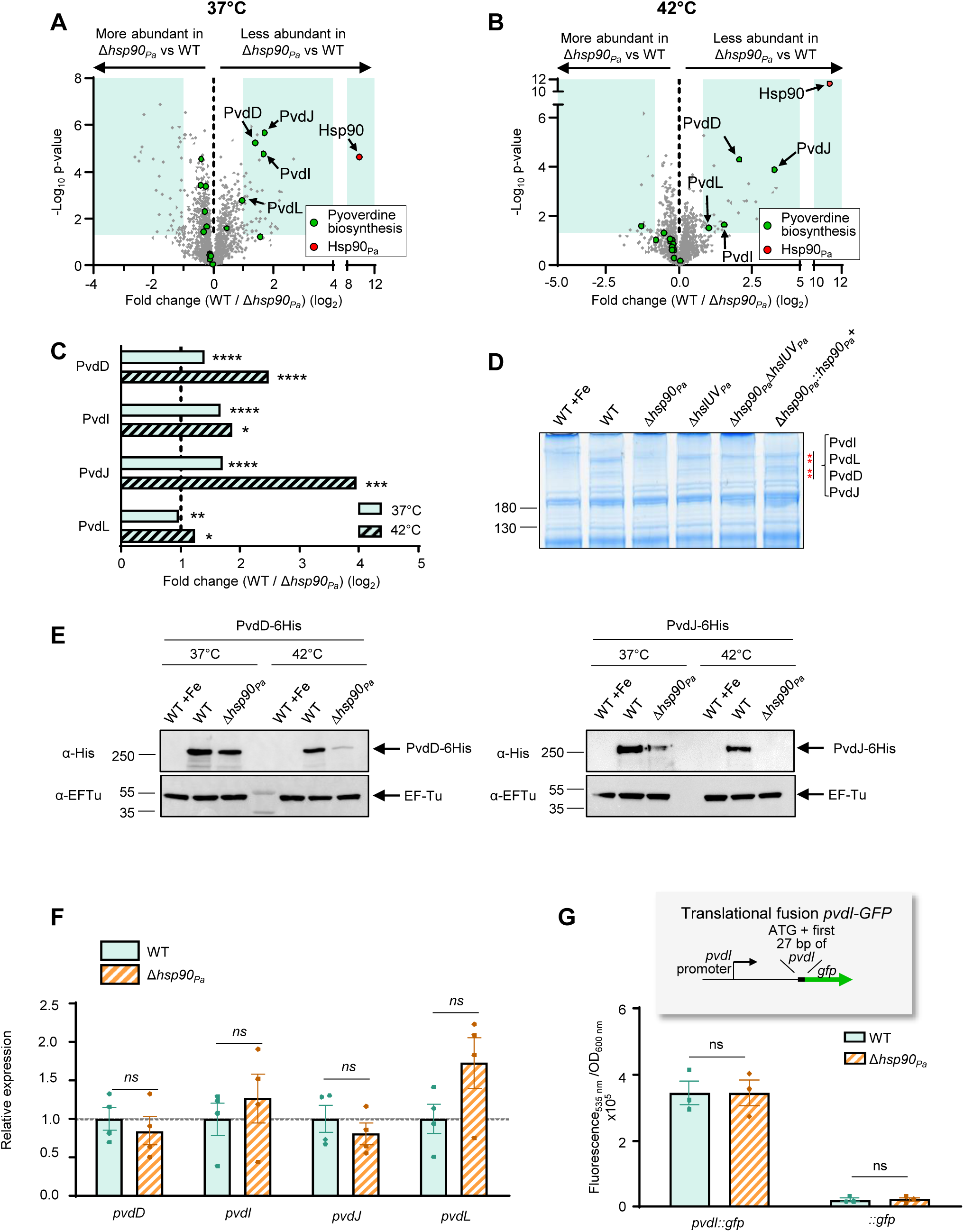
Hsp90*_Pa_* acts at the post-translational level to maintain the abundance of megasynthases involved in pyoverdine production in *P. aeruginosa*. (**A** and **B**) Global comparative quantitative proteomics of *P. aeruginosa* WT and Δ*hsp90_Pa_* strains. The protein content of the two strains grown at 37°C (A) or 42°C (B) in CAA was analyzed by LFQ mass spectrometry, and the relative abundance of proteins is reported as described in Fig. 1B. Hsp90*_Pa_* is colored in red, and proteins involved in pyoverdine biosynthesis in green. The experiments were performed with 5 biological replicates. (C) Relative abundance of megasynthases involved in pyoverdine biosynthesis measured in A and B. p-values associated with each ratio indicate that the differences are significant (****P ≤ 0.0001, ***P ≤ 0.001, **P ≤ 0.01, *P ≤ 0.05). (D) SDS PAGE analysis of high molecular weight proteins. Total protein extracts from WT, Δ*hsp90_Pa_,* Δ*hslUV_Pa_,* Δ*hsp90_Pa_*Δ*hslUV_Pa_* and a complemented strain (Δ*hsp90_Pa_::hsp90_Pa_^+^)* grown in iron depleted medium and iron-supplemented medium (+Fe) at 42°C were separated by SDS-PAGE and stained with Coomassie blue. Mass spectrometry identified PvdD, I, J and L in bands marked with an asterisk. Molecular weight markers (kDa) are indicated on the left of the gel. (E) The amount of PvdD and PvdJ is reduced in the absence of Hsp90*_Pa_*. *P. aeruginosa* WT and Δ*hsp90_Pa_* strains carrying a chromosomal encoded C-terminal His_6_ epitope tagged PvdD (PvdD-6His, 274.5 kDa) or PvdJ (PvdJ-6His, 241 kDa) were grown at 37°C or 42°C in CAA. As a control, the medium was also supplemented with iron (WT+Fe). PvdD or PvdJ was detected by immunoblot with a 6His antibody. EF-Tu (43.3 kDa) was used as loading control. Molecular weight markers (kDa) are indicated on the left. The Western blots shown are representative of two independent experiments. (F) Hsp90*_Pa_* does not modify expression of the genes encoding the megasynthases. mRNA from strains grown at 42°C as in B was extracted, retro-transcribed in cDNA, and quantitative-PCR was performed using specific oligonucleotides to amplify *pvdD*, *pvdI*, *pvdJ*, and *pvdL*. Data are shown as fold variation of expression in the Δ*hsp9*0*_Pa_* strain relative to the WT strain after normalization to *rpoD*, *rpsL*, and 16S rRNA. Dot line (y=1) indicates the relative expression value for which there is no variation compared to the expression in the WT strain. Data from four biological replicates are shown as mean ± SEM. t-test analyses indicate that the differences are not significant (ns). (G) Hsp90*_Pa_* does not modify translation of the Pvd proteins. A 341bp sequence comprising the PvdI promoter (*pvdI*, *pvdJ* and *pvdD* are in operon), RBS, ATG, and the first 27 nucleotides of the *pvdI* coding sequence was cloned upstream of the *gfp* gene and inserted at the *attB* locus on the chromosome of the WT and Δ*hsp90_Pa_* strains. As a control GFP was also inserted at the same site on the chromosome but without any promoter. GFP, whose quantitation reflects the level of translation of PvdI, was measured by fluorescence. Data from three biological replicates are shown as mean ± SEM. t-test analyses indicate that the differences are not significant (ns).

The proteomic results were next confirmed by two additional experiments. First, the WT, Δ*hsp90*_Pa_ and complemented strains were grown in the same conditions as above (CAA minimal medium allowing pyoverdine biosynthesis) but also in iron-replete medium (CAA supplemented with iron as a negative control). The protein content of each strain was separated by SDS-PAGE into a low percentage acrylamide gel and visualized by Coomassie blue as done before to visualize NRPS (53). Four protein bands were observed at high molecular weights (>180 kDa) in the WT strains from iron-depleted but not iron-replete cells (**Fig. 3D**). These protein bands were less intense in the absence of Hsp90_Pa_, and importantly their quantities were restored in the complemented strain. Mass spectrometry identification analysis revealed that the PvdD, PvdI, PvdJ and PvdL proteins were found in this region of the polyacrylamide gel, and they were more abundant in the WT than in the Δ*hsp90*_Pa_ strain, as determined by spectral counting. Second, *pvdD* and *pvdJ* genes were replaced by genes encoding 6His epitope-tagged proteins (PvdD-6His and PvdJ-6His) on the chromosome of the WT and Δ*hsp90*_Pa_ strains. Addition of the 6His tag did not affect the functionality of these proteins (**Fig. S3B**). Strains were grown at 37°C and 42°C in iron-deficient medium, and protein content was separated by SDS-PAGE and analyzed by Western blot with an anti-His antibody. Interestingly, we observed that although the bands corresponding to PvdD-6His and PvdJ-6His were slightly less intense at 37°C in the Δ*hsp90*_Pa_ strains, the abundance of these proteins was dramatically reduced in the absence of Hsp90_Pa_ when strains were grown at 42°C (**Fig. 3E, S3C**). As expected, we found that when iron was added to the culture, PvdD-6His and PvdJ-6His were not produced (58).

Finally, we aimed to rule out a putative regulatory (transcriptional or translational) role of Hsp90_Pa_ in pyoverdine biosynthesis under heat stress rather than a role in NRPS stabilization. We thus performed qRT-PCR experiments (**Fig. 3F**) and used translational GFP fusion to PvdI, the protein encoded by the first gene of the *pvdIJD* operon (**Fig. 3G**) with WT and Δ*hsp90*_Pa_ strains grown at 42°C in iron-deficient medium. These experiments showed on one hand that expression of *pvdD*, *pvdI*, *pvdJ*, and *pvdL* did not significantly vary between the two strains (**Fig. 3F**) and on another hand that GFP levels were not changed between the WT and Δ*hsp90*_Pa_ mutant (**Fig. 3G**) indicating that Hsp90_Pa_ does not play a role in *pvd* genes transcription or PvdI translation

Altogether, these experiments demonstrate that Hsp90_Pa_ is essential to maintain the amount, at the post-translational level, of four NRPS (PvdD, PvdI, PvdJ and PvdL) required for pyoverdine biosynthesis in *P. aeruginosa*. Consequently, a dramatic loss of pyoverdine production is observed in the absence of Hsp90_Pa_ and this phenotype is exacerbated under heat stress.

### The interplay between Hsp90 and the HslUV protease determines the amount of megasynthases

Our results demonstrate that in *E. coli* and *P. aeruginosa*, Hsp90 is required to stabilize megasynthases that are respectively involved in colibactin and pyoverdine biosynthesis. Interestingly, it has previously been shown that in the absence of Hsp90_Ec,_ colibactin production is restored by the deletion of *hslV*_Ec_, the gene encoding the protease component of the chaperone-protease complex HslUV_Ec_ (38). This effect is specific to HslUV, as the absence of another protease, ClpP, does not restore colibactin production (38). This observation suggests that in the absence of Hsp90_Ec_, HslUV_Ec_ degrades one or more yet-to-be discovered Hsp90_Ec_ clients required for colibactin biosynthesis. HslUV is composed of the hexameric ATP-dependent HslU chaperone that unfolds and directly addresses protein substrates to the dodecameric protease component HslV (59–63).

Therefore, we hypothesized that the HslUV_Ec_ protease contributed to the reduced levels of the NRPS ClbH, ClbJ, and ClbN, as well as the hybrid NRPS-PKS ClbB and ClbK, observed in the absence of Hsp90_Ec_. To test for this possibility, three additional comparative quantitative proteomic experiments were performed using two other *E. coli* M1/5 strains in which the *hslV*_Ec_ gene (Δ*hslV*_Ec_), or the *hslV*_Ec_ and *hsp90*_Ec_ genes (Δ*hsp90*_Ec_ Δ*hslV*_Ec_) were deleted (**Fig. 4A, 4B, S4A-C, and Datasets S7 to S12**). Strikingly, we observed that in the double deletion strain (Δ*hsp90*_Ec_ Δ*hslV*_Ec_) the levels of the 5 Clb proteins (ClbB, ClbH, ClbJ, ClbK, and ClbN) which were significantly decreased in the absence of Hsp90_Ec_ were restored or almost restored to the level measured in the WT strain (**Fig. 4A, 4B**, and S4A). The amount of ClbB, ClbH, ClbJ, ClbK, and ClbN was slightly increased in the strain deleted of *hslV*_Ec_ compared to WT, suggesting a small but active turnover of these proteins even in the presence of Hsp90_Ec_ (**Fig. 4B and S4B)**.To confirm that the level of the Clb megasynthases was dependent on HslUV_Ec_, we followed the amount of ClbB with a 6-His epitope tag produced from a plasmid with an arabinose-inducible promoter in the MG1655 *E. coli* strains deleted of *hsp90* and *hslV*. We observed that the amount of ClbB that was strongly reduced in the absence of Hsp90_Ec_ compared to the WT strain as described above, was rescued in the Δ*hsp90*_Ec_Δ*hslV*_Ec_ strain to approximately the level measured in the WT strain (**Fig. 4C, and Fig S4D**). The effect of HslUV_Ec_ was confirmed by complementation experiments in which HslUV_Ec_ was produced from a plasmid (**Fig. S4E**).

**Fig. 4.**
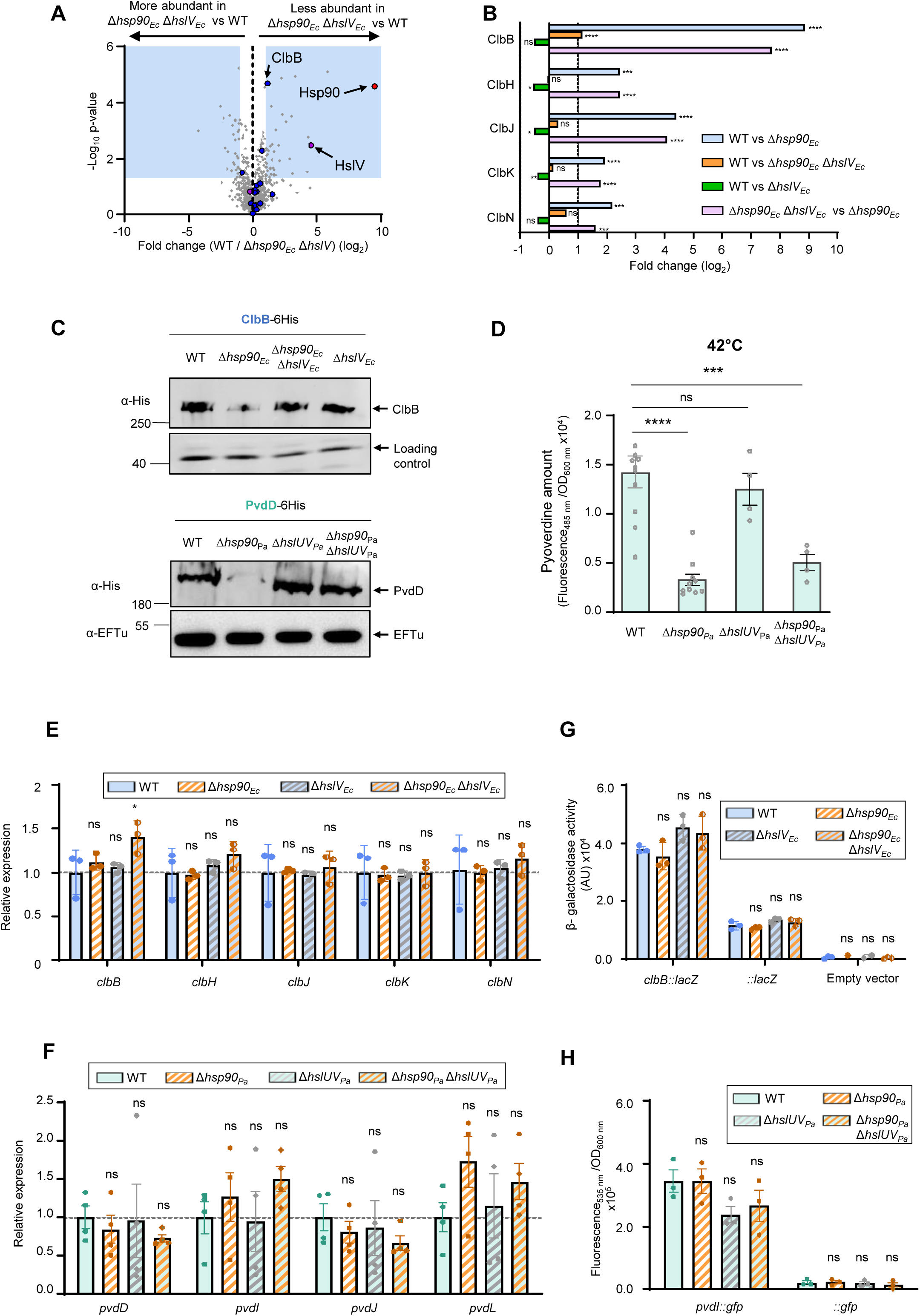
Interplay between Hsp90 and HslUV to control the level of NRPS. (A) Global comparative quantitative proteomics of *E. coli* WT and Δ*hsp90*Δ*hslV* strains. The protein content of the two strains grown at 37°C was analyzed by LFQ mass spectrometry, and the relative abundance of proteins is reported as described in Fig. 1B. Hsp90*_Ec_* is colored in red, and proteins involved in colibactin biosynthesis in blue. The experiments were performed with 4 biological replicates. (B) Relative abundance of megasynthases involved in colibactin biosynthesis measured in Fig. 1B, 4A, and S4A-B. p-values associated with each ratio indicate whether the differences are significant or not (****P ≤ 0.0001, ***P ≤ 0.001, **P ≤ 0.01, *P ≤ 0.05, ns: not significant). (C) The absence of the HslUV protease in the Δ*hsp90* strains increases the level of ClbB in *E. coli* (top panel), or of PvdD in *P. aeruginosa* (bottom panel). (Top panel) *E. coli* strains carrying a plasmid allowing the production of ClbB-6His (353.4 kDa) were grown at 37°C and arabinose was added to induce ClbB-6His production. ClbB-6His was detected using a 6His antibody. The band labelled ‘‘loading control’’ corresponds to a protein that is nonspecifically detected by the 6His antibody. (Bottom panel) *P. aeruginosa* strains carrying a chromosomal encoded C-terminal 6-His epitope tagged PvdD (PvdD-6His, 274.5 kDa) were grown at 42°C in CAA. PvdD was detected by immunoblot with a 6His antibody. Antibody against EF-Tu (43.3 kDa) was used as loading control. Molecular weight markers (kDa) are indicated on the left. The Western blots shown are representative of three independent experiments. (D) Pyoverdine production analysis. *P. aeruginosa* WT, Δ*hsp90*, Δ*hslUV*, and Δ*hsp90* Δ*hslUV* strains were grown at 42°C in CAA. After 5h growth, pyoverdine was quantified by measuring fluorescence (excitation: 400 nm; emission: 485 nm) that was standardized to the number of cells (OD_600_). Data from at least four replicates are shown as mean ± SEM. (**E**, **F**) The absence of the HslUV protease does not affect the expression of the genes encoding the megasynthases. qRT-PCR experiments were performed on (E) the *E. coli* strains Δ*hslV_Ec_* and Δ*hsp90_Ec_* Δ*hslV_Ec_* prepared as in Fig. 1D, and on (F) the *P. aeruginosa* strains Δ*hslUV_Pa_* and Δ*hsp90_Pa_* Δ*hslUV_Pa_* as in Fig. 3F. Note that the controls shown here for WT and Δ*hsp90* strains have been presented in Fig. 1D and in Fig. 3F for *E. coli* and *P. aeruginosa* strains, respectively. Data from three (E) or four (F) biological replicates are shown as mean ± SEM. (**G**, **H**) The absence of the HslUV protease does not affect translation of the Clb or Pvd proteins. (G) The plasmid allowing measurement of the *clbB::lacZ* translational fusion was introduced into the *E. coli* strains Δ*hslV_Ec_* and Δ*hsp90_Ec_* Δ*hslV_Ec_*, and the activity of the fusion was measured as shown in Fig. 1E. (H) Chromosomal translational GFP fusion to PvdI was inserted into the chromosome of *P. aeruginosa* in the Δ*hslUV_Pa_* and Δ*hsp90_Pa_* Δ*hslUV_Pa_* strains. GFP, whose quantitation reflects the level of translation of PvdI, was measured by fluorescence as in Fig. 3G. Note that the controls shown here for WT and Δ*hsp90* strains have been presented in Fig. 1E and in Fig. 3G for *E. coli* and *P. aeruginosa* strains, respectively. Data from three biological replicates are shown as mean ± SEM. In D-H, results of one-way ANOVA tests indicate whether the differences measured are significant (*: p-value ≤ 0.05; ***: p-value ≤ 0.001; ****: p-value ≤ 0.0001;) or not significant (ns, p-value > 0.05).

Next, we asked whether the NRPS involved in pyoverdine biosynthesis in *P. aeruginosa* were also reduced in a HslUV_Pa_-dependent manner in the absence of Hsp90_Pa_. To this end, a strain deleted for *hslUV*_Pa_ and a triple mutant Δ*hsp90*_Pa_Δ*hslUV*_Pa_ were constructed. The strains were grown in minimal medium at 42°C and protein content was analyzed by Coomassie blue stained SDS-PAGE (**Fig. 3D**). Interestingly, the four protein bands whose intensity was dramatically reduced in the absence of Hsp90_Pa_ were restored in the Δ*hsp90*_Pa_Δ*hslUV*_Pa_ strain to a level similar to the one observed in the WT strain (**Fig. 3D**). Then, a sequence encoding a 6-His tag was introduced at the 3’-end of the *pvdD* or *pvdJ* gene in these strains. After growth of the strains producing PvdD-6His or PvdJ-6His at 42°C in minimal medium, total protein extracts were separated on SDS-PAGE, and PvdD or PvdJ was detected on Western blot with an anti-His antibody. Remarkably, deleting *hslUV*_Pa_ in the Δ*hsp90*_Pa_ background resulted in a dramatic increase in the amount of PvdD and PvdJ compared to the Δ*hsp90*_Pa_ mutant and a restoration even above the WT level (**Fig. 4C and S4D and S4F**). These results were confirmed by complementation experiments in which *hslUV* was introduced into the chromosome of the *P. aeruginosa* strain Δ*hsp90*_Pa_ Δ*hslUV*_Pa_ (**Fig. S4G**). To assess if restoration of PvdD and PvdJ levels translates into pyoverdine biosynthesis, we next measured pyoverdine production in the various strains. While pyoverdine level was similar to a WT strain in the Δ*hslUV*_Pa_ mutant, it was only slightly increased in the Δ*hsp90*_Pa_Δ*hslUV*_Pa_ compared to the Δ*hsp90*_Pa_ strain (**Fig. 4D and S4H**). These results are in agreement with a model in which in stress condition (42°C) Hsp90_Pa_ not only stabilizes the Pvd proteins, but also acts in their folding or activation.

Finally, to exclude the possibility that the restoration of the megasynthase levels observed in the absence of HslUV in the *hsp90* mutant background of both bacteria was due to increased transcription or translation, qRT-PCR and translational fusions were used in these strains as in Fig. **1D-E** and **3F-G**. We found that the absence of the HslUV protease had little or no effect on the expression of the genes encoding the megasynthases, compared to the WT and *hsp90* mutant strains in both *E. coli* and *P. aeruginosa* (**Fig. 4E and 4F**). Similarly, no significant variation was measured among the four strains with the translational fusions (**Fig. 4G and 4H**).

Al together, these experiments show that Hsp90 prevents, at the post-translational level, the HslUV-dependent decrease of megasynthase abundance.

### Hsp90 is required to maintain the levels of additional NRPS and PKS-NRPS enzymes

Our results indicate that biosynthesis of colibactin in *E. coli* and pyoverdine in *P. aeruginosa* is dramatically reduced in the absence of Hsp90 due to a decreased abundance of large enzymes that belong to the NRPS family (NRPS only: ClbH, J, N for colibactin, and PvdD, I, J, and L for pyoverdine; hybrid NRPS-PKS: ClbB and K for colibactin) (**Fig. 5A**).

**Fig. 5.**
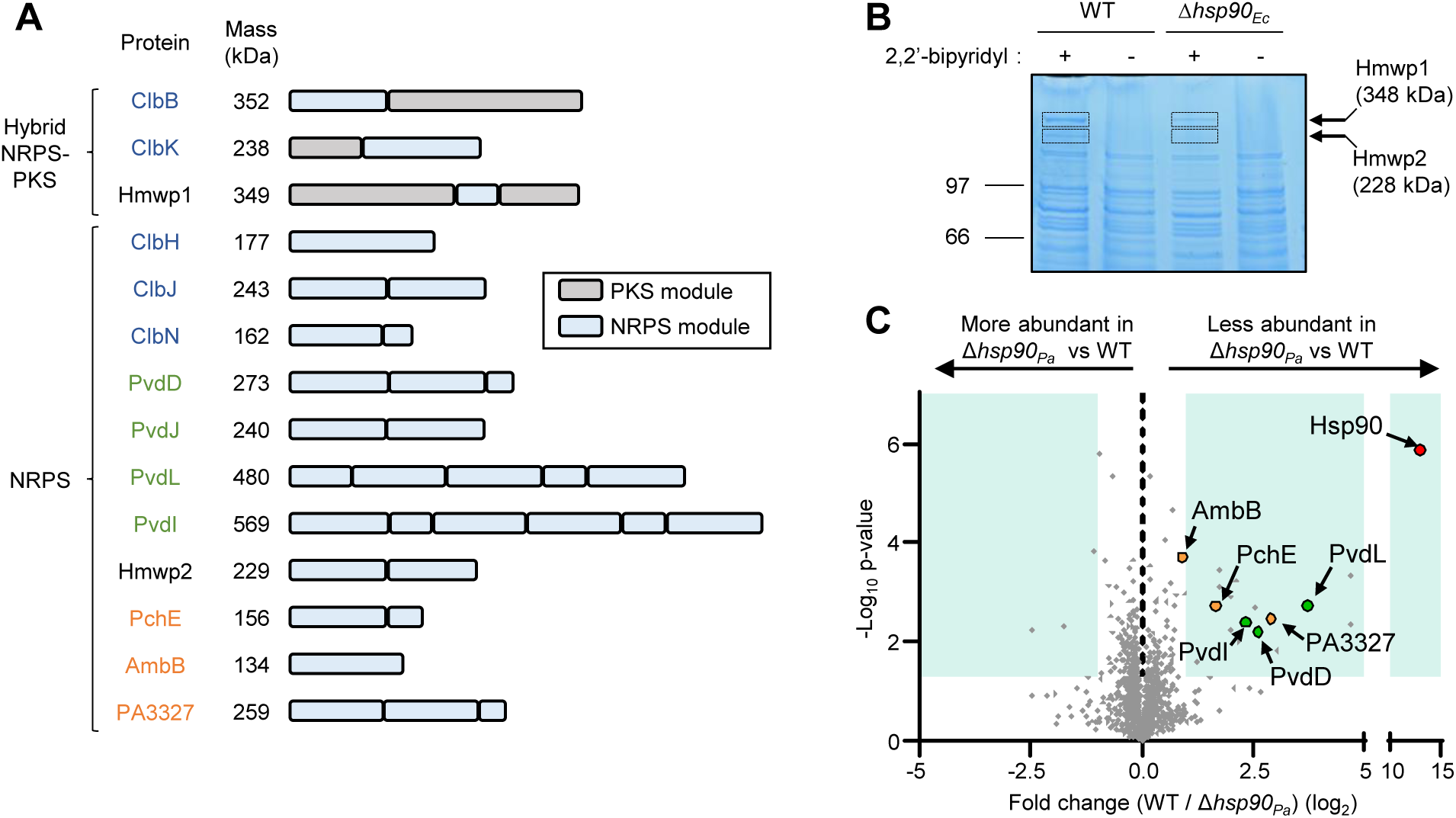
The levels of various NRPS megaenzymes depend on the Hsp90 chaperone. (A) Schematic representation of the modular organization of the megasynthases containing NRPS modules whose amount is reduced in the absence of Hsp90. Shown are the hybrid PKS-NRPS ClbB, ClbK, and Hmwp1 from *E. coli*, and the NRPS ClbH, ClbJ, ClbN, and Hmwp2 from *E. coli*, and PvdD, PvdJ, PvdL, PvdI, PchE, AmbB and PA3327 from *P. aeruginosa*. The molecular mass of the proteins is indicated. Proteins labeled in blue are involved in the biosynthesis of colibactin, in green of pyoverdine, and in black of yersiniabactin. In orange, PchE, AmbB and PA3327 are involved in the biosynthesis of pyochelin, AMB and azetidomonamide A, respectively. (**B**) Megasynthases involved in yersiniabactin biosynthesis are less abundant in the absence of Hsp90*_Ec_*. *E. coli* WT and Δ*hsp90* strains were grown in LB rich media at 37°C in the presence or not of the iron chelator bipyridyl (200 µM). The same number of cells was collected, resuspended in denaturing loading buffer, heat-treated at 95°C, and proteins were separated by SDS-PAGE before to be stained with Coomassie blue. The protein content of the framed bands has been identified by mass spectrometry, and they mainly contain HMWP1 (upper band) or HMWP2 (lower band) as indicated by the arrows. The molecular weight markers (kDa) are indicated on the left of the gel. The gel shown is representative of three independent experiments, and three independent mass spectrometry analysis were performed. (**C**) Global comparative quantitative proteomics of *P. aeruginosa* WT and Δ*hsp90* strains. The protein content of the two strains grown in LB rich medium at 37°C was analyzed by LFQ mass spectrometry, and the relative abundance of proteins is reported as described in Fig. 1B. Hsp90*_Pa_* is colored in red, NRPS involved in pyoverdine biosynthesis in green, and additional NRPS proteins are colored orange. The experiments were performed with 3 biological replicates.

Hsp90 has also previously been described in the literature to be required for the production of other NRP or NRP-PK compounds including the yersiniabactin siderophore, without a molecular explanation on how this occurs (38, 42, 43). Interestingly, Hmwp1, a hybrid NRPS-PKS and one of the two megasynthases involved in yersiniabactin biosynthesis, was less abundant in the absence of Hsp90_Ec_ in the global comparative proteomics analysis presented above, although the growth conditions (rich media) did not favour production of yersiniabactin (**Dataset S2**). To confirm this data and with the idea of generalizing our model of megasynthase control by Hsp90, we explored the biosynthesis of the yersiniabactin siderophore in *E. coli*. Yersiniabactin production requires more than 10 proteins including two megasynthases, the hybrid NRPS-PKS Hmwp1 and the NRPS Hmwp2 (39). In order to visualize Hmwp1 and Hmwp2 in *E. coli* wild-type and Δ*hsp90,* strains were grown in the presence or absence of the iron chelator 2,2’-bipyridyl, since iron deficiency increases yersiniabactin production (39, 64, 65), and total protein extracts were separated on SDS-PAGE. The high molecular weight of Hmwp1 and Hmwp2, 348 kDa and 228 kDa, respectively, allowed their visualization on Coomassie blue stained gel. When grown in the presence of 2,2’-bipyridyl (i.e. iron deficiency), two high molecular weight bands were present in the WT strain (**Fig. 5B**) whereas they were not visible without 2,2’-bipyridyl. Mass spectrometry analysis identified these as Hmwp1 and Hmwp2. Interestingly, we observed that the intensity of these bands was strongly decreased in the absence of Hsp90_Ec_. These results indicate that, as for colibactin and pyoverdine biosynthesis, Hsp90_Ec_ leads to higher levels of the Hmwp1 and Hmwp2 megasynthases to allow yersiniabactin production.

To search for other secondary metabolites synthesized by NRP enzymes, we inspected the *P. aeruginosa* genome using the antiSMASH software (https://antismash.secondarymetabolites.org). In addition to pyoverdine, three additional secondary metabolites synthesized by NRPS were identified: pyochelin, azetidomonamide A and L-2-amino-4-methoxy-trans-3-butenoic acid (AMB) (**Fig. 5A**). Strikingly, global proteomics revealed that Hsp90 mutation had an impact on three NRPS involved in the biosynthesis of these secondary metabolites. Indeed, a comparison of the proteome of *P. aeruginosa* WT and Δ*hsp90* strains grown in rich media at 37°C revealed that the level of PchE, AmbB and PA3327 involved in the biosynthesis of pyochelin, AMB and azetidomonamide A, respectively, were reduced in the absence of Hsp90_Pa_ (**Fig. 5C, Datasets S13 and S14**) (66–69). These results suggest that Hsp90_Pa_ is involved in all NRPS-dependent secondary metabolite biosynthesis encoded on the *P. aeruginosa* genome.

Together, these results show that the role of Hsp90 in maintaining a proper level of megasynthases required for NRP or PK-NRP compound production is widespread among several biosynthetic pathways not only including colibactin and pyoverdine, but also yersiniabactin, pyochelin, and others.

## Discussion

How Hsp90 participates in bacterial physiology and virulence has been elusive for many years. Here, we demonstrate that one crucial role of bacterial Hsp90 is to maintain the right amount of large enzymes (megasynthases) required to produce secondary metabolites that belong to the classes of non-ribosomal peptide or hybrid polyketide non-ribosomal peptide. Indeed, we found that the level of 5 megasynthases involved in colibactin biosynthesis in *E. coli* (ClbB, ClbJ, ClbH, ClbN, and ClbK) (**Fig. 1**), and 4 megasynthases involved in pyoverdine biosynthesis in *P. aeruginosa* (PvdD, PvdI, PvdJ, and PvdL) (**Fig. 3**) was dramatically reduced in the absence of Hsp90. qRT-PCR and translational fusions strongly suggest that Hsp90 controls the levels of megasynthases at the post-translational level (**Fig. 1 and 3**), and we showed an interplay between Hsp90 and a protease, HslUV, in regulating megasynthase abundance (**Fig. 4**). We also identified several additional megasynthases (Hmwp1, Hmwp2, PchE, AmbB, and PA3327) in *E. coli* and *P. aeruginosa* whose levels were reduced in the absence of Hsp90 (**Fig. 5**), suggesting a general role of Hsp90 in maintaining the abundance of this kind of enzymes. In addition to establishing the role of Hsp90 in controlling the amount of NRPS and PKS-NRPS protein, our study provides a large set of comparative proteomic data with information on the impact of Hsp90 on the proteome of *E. coli* grown at 37°C (**Datasets S1 and S2**), and of *P. aeruginosa* grown both at 37°C and 42°C in minimal medium (**Datasets S3 to S6**), and in rich medium at 37°C (**Datasets S13 and S14**). Proteomic analyses of various *E. coli* strains deleted of the HslV protease are also presented (**Datasets S7 and S12**). Data from these proteomic analyses are under study and will be exploited in future work. Interestingly, the megasynthases whose quantity is controlled by Hsp90 either belong to the NRPS or to the hybrid PKS-NRPS enzyme families, strongly suggesting that the NRPS moiety of the hybrid proteins is recognized by Hsp90. Two other observations reinforce this idea: the levels of the PKS enzymes required for colibactin production (ClbC, ClbI, and ClbO) were not reduced in the absence of Hsp90_Ec_ (**Fig. S1A**), and Hsp90 is required for the biosynthesis of pyoverdine and arthrofactin that are NRP-only molecules (43). To our knowledge a role of bacterial Hsp90 in the biosynthesis of PKS molecules has not been reported yet. Our results thus suggest that NRP and hybrid PK-NRP megasynthases are clients of Hsp90. Future work will focus on the investigation of the interaction between Hsp90 and these large enzymes. They are composed of one or several modules of three essential conserved domains that together allow the assembly of the nascent molecule: a condensation domain, an adenylation domain, and a carrier domain. Alphafold3 structure predictions showed a strong overlap (RMSD between 0.7 and 1.4 angstroms on pruned atom pairs) for each NRPS domain of PvdD, PvdJ, ClbB, and ClbJ (**Fig. S5**), suggesting that a common fold or domain that remains to be discovered could be recognized by Hsp90. It is also possible that Hsp90 recognizes non-native or transient folding intermediates rather than fully folded mature conformations which will make identifying its recognition features more challenging.

These observations raise the question of whether Hsp90 controls the abundance of all NRPS enzymes or whether its role is specific to some of them. Garcie *et al*. have observed that the production of enterobactin and salmochelin, two NRP metabolites, does not vary in the absence of Hsp90 (38). Similarly, we identified NRPS proteins in the proteomic analysis (including for example PchF, which is involved in pyochelin biosynthesis) whose amount was not affected by the absence of Hsp90_Pa_ under our experimental conditions. These examples may indicate that some NRPS are clients of Hsp90 while others are not, due to some yet unknown characteristics. However, it is important to note that the role of Hsp90_Pa_ in maintaining the level of NRPS required for pyoverdine production depended on the temperature, with a greater Hsp90_Pa_ requirement for PvdD, PvdI, PvdJ and PvdL at 42°C than at 37°C (**Fig. 3C**). Interestingly, this correlated with the amount of pyoverdine produced since Hsp90_Pa_ was crucial for pyoverdine production under heat stress (42°C) whereas the Δ*hsp90*_Pa_ mutant strain produced the same level of pyoverdine as the WT at 30°C (**Fig. 2 and S2B**). Finally, the abundance of PchE and PA3327 which did not vary in minimal medium between the WT and Δ*hsp90*_Pa_ strains was decreased in the absence of Hsp90_Pa_ in rich medium (**Fig. 5C**). Altogether these results suggest that Hsp90 is essential for the production of most of the NRP or PK-NRP molecules under specific conditions which may be difficult to identify. In contrast, for some systems such as colibactin biosynthesis, Hsp90 is essential even under non-stressed conditions (37°C) (38).

The interplay we observed between Hsp90 and HslUV in maintaining megasynthase abundance, together with our finding that the transcription and translation of the *pvd* and *clb* genes encoding these megasynthases are unaffected by Hsp90 and/or HslUV (**Fig. 4**), supports a model in which HslUV directly degrades the megasynthases in the absence of Hsp90. This model of protection and folding by Hsp90, and degradation by the protease HslUV has already been proposed (38, 49, 70). For example, TilS, an Hsp90 client in *S. oneidensis* is degraded by HslUV in the absence of Hsp90 (33, 49). Since TilS is an essential protein, *S. oneidensis* without Hsp90 cannot grow due to the absence of TilS. Deletion of *hslUV* in the Δ*hsp90* background suppresses the growth defect of the Δ*hsp90* strain. A similar result was obtained when measuring colibactin production, whose amount strongly decreased in an *E. coli* Δ*hsp90* mutant, and was restored in a Δ*hsp90*Δ*hslV* strain (38). Our results provide the molecular explanation for this phenotype since, we found that the levels of the 5 NRPSs that were reduced in the *E. coli* Δ*hsp90* strain were restored to WT level when the protease was inactivated in the *hsp90*_Ec_ deletion background (**Fig. 4B**). These results suggest that the main role of Hsp90_Ec_ in colibactin production under non-stress conditions (37°C) is to protect these enzymes from degradation, since blocking their degradation in the absence of Hsp90_Ec_ is sufficient to restore colibactin production. Interestingly, an alternative model can be proposed with pyoverdine biosynthesis. We found that under heat stress (42°C), although the amount of the NRP enzymes was even higher in the Δ*hsp90*_Pa_Δ*hslUV*_Pa_ strain compared to the level measured in the WT strain (**Fig. 4C, S4D, and S4F**), pyoverdine production was poorly restored in the Δ*hsp90*_Pa_ Δ*hslUV*_Pa_ strain compared to the Δ*hsp90*_Pa_ strain and did not reach the WT level (**Fig. 4D**). Therefore, our results suggest that under heat stress, the role of Hsp90 in *P. aeruginosa* cannot be limited to the protection against degradation of NRPS, and Hsp90_Pa_ most probably is also actively required for the folding or activation of these enzymes.

Our study adds a new level of understanding and complexity to the biosynthesis of the NRP and PK-NRP secondary metabolites by demonstrating the role and importance of the Hsp90 chaperone in this process. We have previously shown that Hsp90 cooperates with the DnaK chaperone in the production of colibactin (71). A similar chaperone pathway may exist for the biosynthesis of other NRP and PK-NRP molecules. Based on several studies from different laboratories, the Hsp70/DnaK chaperone first interacts with clients, and transfers them to Hsp90 to allow final folding or activation (14, 17, 72). If the DnaK-Hsp90 cycle fails, some proteases, such as here HslUV, degrade the clients.

Importantly our work demonstrates a key role for Hsp90 in maintaining the abundance of NRP and PK-NRP megasynthases involved in the biosynthesis of multiple virulence factors critical for *E. coli* and *P. aeruginosa* pathogenicity. For example, during infection, the host immune system sequesters trace elements, such as iron to limit pathogenicity in a process called nutritional immunity (73). In the case of iron, bacteria produce in turn low molecular weight molecules called siderophores, which are secreted, and interact with iron before being internalized into the bacteria to increase intracellular iron concentrations and counteract iron limitation (52). We show that the biosynthesis of at least two siderophores, yersiniabactin in *E. coli* and pyoverdine in *P. aeruginosa*, and possibly also pyochelin in *P. aeruginosa* (via PchE) is impacted by the absence of Hsp90. Hsp90 is also involved in the biosynthesis of bacterial toxins that are important weapons that bacteria use to damage host cells during infection. It is required for the biosynthesis of colibactin, a toxin produced by *E. coli* that causes DNA damage and mutagenesis in bacterial and eukaryotic host cells, and albicidin, a toxin that blocks sugarcane chloroplast differentiation in *Xanthomonas albilineans* (41, 42). Hsp90 could also participate in the production of the toxin AMB (L-2-amino-4-methoxy-trans-3-butenoic acid) in *P. aeruginosa* by maintaining AmbB amount, although this remains to be tested. In addition, by stabilizing the NRPS encoded by the PA3327 gene, which produces azetidomonamide A in *P. aeruginosa*, Hsp90 could have a broad effect on pathogenicity and virulence as this protein has previously been implicated in biofilm formation and in phenazine and alkyl quinolone production (69). Finally, Hsp90 may also be involved in bacterial motility by protecting an unidentified megasynthase to produce the biosurfactant arthrofactin in *Pseudomonas* sp. MIS38 (43). Given the general role of Hsp90 in NRPS stabilization that we have uncovered, one can speculate that Hsp90 is likely to participate in the biosynthesis of a wide range of virulence factors in pathogenic bacteria.Our study therefore provides a strong rationale and molecular evidence for considering bacterial Hsp90 as a promising target for anti-virulence strategies aimed at disarming pathogenic bacteria. Finally a better understanding of how proteins responsible for the production of virulence factors are synthesized, protected, folded and/or degraded is crucial for developing new methods to inhibit bacterial virulence. In addition, the controlled use of chaperones or the inhibition of certain proteases could improve the production of NRP and PK-NRP molecules for biotechnological purposes.

## Materials and Methods

### Strains, plasmids, and growth conditions

Strains, plasmids, and primers used in this study are listed in the **Supporting Tables S1, S2, and S3**, respectively. The wild-type commensal *E. coli* strain used for colibactin production is M1/5 (45). The wild-type *P. aeruginosa* strain is PAO1. Construction of strains and plasmids, and growth conditions are described in Supporting Methods.

### Assays

For proteomic analysis, samples were prepared as described (74) with some modifications (Supporting Methods). Other assays including qRT-PCR, translational fusions, extracellular pyoverdine measurement, as well as detection of ClbB, the Pvd proteins, and the proteins involved in yersiniabactin production are described in Supporting Methods.

### Data availability

The mass spectrometry proteomics data have been deposited to the ProteomeXchange Consortium via the PRIDE (75) partner repository with the dataset identifier PXD056877.

### Statistical analysis

Statistical analysis was performed using the GraphPad Prism 10.2.3 software. One-way analysis of variance followed by Dunnett’s post-hoc test (multiple comparisons) was used for comparisons between three or more groups, and the unpaired t-test (single comparison) was used for comparisons between two groups.

## Supporting information

Supporting Materials and Methods; Figures S1 to S5; Tables S1 to S4; SI References

## Acknowledgments

We thank members of our groups for their help and fruitful discussions, and especially Elodie Bergé for her precious contribution. We thank Yann Denis from the transcriptomic platform of the IMM (CNRS) for assistance with cDNA preparation and qPCR, and Eliot David for his support in some of our experiments. A.F is supported by a PhD studentship from AMU. This work was supported by the CNRS, AMU, INSERM, INRAE, ENVT, Université Paul Sabatier, and ANR (ANR-20-CE44-0017 and ANR-23-CE44-0018). The Marseille Proteomics platform (MaP; marseille-proteomique.univ-amu.fr) is IBiSA- and AMU labeled technological platform. The authors declare no competing interests.

## References

1. I. Matic, Mutation Rate Heterogeneity Increases Odds of Survival in Unpredictable Environments. Mol. Cell 75, 421–425 (2019).

2. E. A. Mueller, P. A. Levin, Bacterial Cell Wall Quality Control during Environmental Stress. mBio 11, e02456–20 (2020).

3. Y. Zhang, C. A. Gross, Cold Shock Response in Bacteria. Annu. Rev. Genet. 55, 377–400 (2021).

4. R. Njenga, J. Boele, Y. Öztürk, H.-G. Koch, Coping with stress: How bacteria fine-tune protein synthesis and protein transport. J. Biol. Chem. 299, 105163 (2023).

5. K. Schumacher, S. Brameyer, K. Jung, Bacterial acid stress response: from cellular changes to antibiotic tolerance and phenotypic heterogeneity. Curr. Opin. Microbiol. 75, 102367 (2023).

6. S. Bouillet, T. S. Bauer, S. Gottesman, RpoS and the bacterial general stress response. Microbiol. Mol. Biol. Rev. MMBR 88, e0015122 (2024).

7. F. C. Fang, E. R. Frawley, T. Tapscott, A. Vázquez-Torres, Bacterial Stress Responses during Host Infection. Cell Host Microbe 20, 133–143 (2016).

8. K. G. K. Goh, et al., An opportunistic pathogen under stress: how Group B Streptococcus responds to cytotoxic reactive species and conditions of metal ion imbalance to survive. FEMS Microbiol. Rev. 48, fuae009 (2024).

9. P. Genevaux, C. Georgopoulos, W. L. Kelley, The Hsp70 chaperone machines of Escherichia coli: a paradigm for the repartition of chaperone functions. Mol. Microbiol. 66, 840–857 (2007).

10. D. Balchin, M. Hayer-Hartl, F. U. Hartl, In vivo aspects of protein folding and quality control. Science 353, aac4354 (2016).

11. V. Dahiya, J. Buchner, Functional principles and regulation of molecular chaperones. Adv. Protein Chem. Struct. Biol. 114, 1–60 (2019).

12. R. Rosenzweig, N. B. Nillegoda, M. P. Mayer, B. Bukau, The Hsp70 chaperone network. Nat. Rev. Mol. Cell Biol. 20, 665–680 (2019).

13. S. Wickner, T.-L. L. Nguyen, O. Genest, The Bacterial Hsp90 Chaperone: Cellular Functions and Mechanism of Action. Annu. Rev. Microbiol. 75, 719–739 (2021).

14. A. C. Wickramaratne, S. Wickner, A. N. Kravats, Hsp90, a team player in protein quality control and the stress response in bacteria. Microbiol. Mol. Biol. Rev. MMBR 88, e0017622 (2024).

15. F. H. Schopf, M. M. Biebl, J. Buchner, The HSP90 chaperone machinery. Nat. Rev. Mol. Cell Biol. 18, 345–360 (2017).

16. M. M. Biebl, J. Buchner, Structure, Function, and Regulation of the Hsp90 Machinery. Cold Spring Harb. Perspect. Biol. 11 (2019).

17. O. Genest, S. Wickner, S. M. Doyle, Hsp90 and Hsp70 chaperones: Collaborators in protein remodeling. J. Biol. Chem. 294, 2109–2120 (2019).

18. G. Chiosis, C. S. Digwal, J. B. Trepel, L. Neckers, Structural and functional complexity of HSP90 in cellular homeostasis and disease. Nat. Rev. Mol. Cell Biol. 24, 797–815 (2023).

19. M. M. U. Ali, et al., Crystal structure of an Hsp90-nucleotide-p23/Sba1 closed chaperone complex. Nature 440, 1013–1017 (2006).

20. A. K. Shiau, S. F. Harris, D. R. Southworth, D. A. Agard, Structural Analysis of E. coli hsp90 reveals dramatic nucleotide-dependent conformational rearrangements. Cell 127, 329–340 (2006).

21. D. E. Dollins, J. J. Warren, R. M. Immormino, D. T. Gewirth, Structures of GRP94-nucleotide complexes reveal mechanistic differences between the hsp90 chaperones. Mol. Cell 28, 41– 56 (2007).

22. M. P. Mayer, Gymnastics of molecular chaperones. Mol. Cell 39, 321–331 (2010).

23. K. A. Krukenberg, T. O. Street, L. A. Lavery, D. A. Agard, Conformational dynamics of the molecular chaperone Hsp90. Q. Rev. Biophys. 44, 229–255 (2011).

24. D. R. Southworth, D. A. Agard, Species-dependent ensembles of conserved conformational states define the Hsp90 chaperone ATPase cycle. Mol. Cell 32, 631–640 (2008).

25. J. L. Johnson, Evolution and function of diverse Hsp90 homologs and cochaperone proteins. Biochim. Biophys. Acta 1823, 607–613 (2012).

26. J. Li, J. Soroka, J. Buchner, The Hsp90 chaperone machinery: conformational dynamics and regulation by co-chaperones. Biochim. Biophys. Acta 1823, 624–635 (2012).

27. Y. Miyata, H. Nakamoto, L. Neckers, The therapeutic target Hsp90 and cancer hallmarks. Curr. Pharm. Des. 19, 347–365 (2013).

28. L. M. Butler, R. Ferraldeschi, H. K. Armstrong, M. M. Centenera, P. Workman, Maximizing the Therapeutic Potential of HSP90 Inhibitors. Mol. Cancer Res. MCR 13, 1445–1451 (2015).

29. S. Rastogi, et al., An update on the status of HSP90 inhibitors in cancer clinical trials. Cell Stress Chaperones 29, 519–539 (2024).

30. N. Tanaka, H. Nakamoto, HtpG is essential for the thermal stress management in cyanobacteria. FEBS Lett. 458, 117–123 (1999).

31. M. M. Hossain, H. Nakamoto, HtpG plays a role in cold acclimation in cyanobacteria. Curr. Microbiol. 44, 291–296 (2002).

32. M. M. Hossain, H. Nakamoto, Role for the cyanobacterial HtpG in protection from oxidative stress. Curr. Microbiol. 46, 70–76 (2003).

33. F. A. Honoré, V. Méjean, O. Genest, Hsp90 Is Essential under Heat Stress in the Bacterium Shewanella oneidensis. Cell Rep. 19, 680–687 (2017).

34. J. C. Bardwell, E. A. Craig, Ancient heat shock gene is dispensable. J. Bacteriol. 170, 2977– 2983 (1988).

35. E. Verbrugghe, et al., HtpG contributes to Salmonella Typhimurium intestinal persistence in pigs. Vet. Res. 46, 118 (2015).

36. A. M. King, et al., High-temperature protein G is an essential virulence factor of Leptospira interrogans. Infect. Immun. 82, 1123–1131 (2014).

37. W. Dang, Y. Hu, L. Sun, HtpG is involved in the pathogenesis of Edwardsiella tarda. Vet. Microbiol. 152, 394–400 (2011).

38. C. Garcie, et al., The Bacterial Stress-Responsive Hsp90 Chaperone (HtpG) Is Required for the Production of the Genotoxin Colibactin and the Siderophore Yersiniabactin in Escherichia coli. J. Infect. Dis. 214, 916–924 (2016).

39. R. D. Perry, J. D. Fetherston, Yersiniabactin iron uptake: mechanisms and role in Yersinia pestis pathogenesis. Microbes Infect. 13, 808–817 (2011).

40. F. Taieb, C. Petit, J.-P. Nougayrède, E. Oswald, The Enterobacterial Genotoxins: Cytolethal Distending Toxin and Colibactin. EcoSal Plus 7 (2016).

41. C. V. Chagneau, et al., The pks island: a bacterial Swiss army knife?: Colibactin: beyond DNA damage and cancer. Trends Microbiol. S0966–842X(22)00122–6 (2022). 10.1016/j.tim.2022.05.010.

42. E. Vivien, et al., Xanthomonas albilineans HtpG is required for biosynthesis of the antibiotic and phytotoxin albicidin. FEMS Microbiol. Lett. 251, 81–89 (2005).

43. K. Washio, S. P. Lim, N. Roongsawang, M. Morikawa, Identification and characterization of the genes responsible for the production of the cyclic lipopeptide arthrofactin by Pseudomonas sp. MIS38. *Biosci. Biotechnol. Biochem.* **74**, 992–999 (2010).

44. A. M. Grudniak, B. Klecha, K. I. Wolska, Effects of null mutation of the heat-shock gene htpG on the production of virulence factors by Pseudomonas aeruginosa. Future Microbiol. 13, 69– 80 (2018).

45. A. Wallenstein, et al., ClbR Is the Key Transcriptional Activator of Colibactin Gene Expression in Escherichia coli. mSphere 5, e00591–20 (2020).

46. J.-P. Nougayrède, et al., Escherichia coli induces DNA double-strand breaks in eukaryotic cells. Science 313, 848–851 (2006).

47. M. Xue, et al., Structure elucidation of colibactin and its DNA cross-links. Science 365, eaax2685 (2019).

48. E. Addington, S. Sandalli, A. J. Roe, Current understandings of colibactin regulation. Microbiol. Read. Engl. 170, 001427 (2024).

49. F. A. Honoré, N. J. Maillot, V. Méjean, O. Genest, Interplay between the Hsp90 Chaperone and the HslVU Protease To Regulate the Level of an Essential Protein in Shewanella oneidensis. mBio 10, e00269–19 (2019).

50. A. Balasubramanian, M. Markovski, J. R. Hoskins, S. M. Doyle, S. Wickner, Hsp90 of E. coli modulates assembly of FtsZ, the bacterial tubulin homolog. Proc. Natl. Acad. Sci. U. S. A. 116, 12285–12294 (2019).

51. M. T. Ringel, T. Brüser, The biosynthesis of pyoverdines. Microb. Cell Graz Austria 5, 424– 437 (2018).

52. I. J. Schalk, Bacterial siderophores: diversity, uptake pathways and applications. Nat. Rev. Microbiol. (2024). 10.1038/s41579-024-01090-6.

53. F. Imperi, P. Visca, Subcellular localization of the pyoverdine biogenesis machinery of Pseudomonas aeruginosa: a membrane-associated “siderosome.” FEBS Lett. 587, 3387– 3391 (2013).

54. V. Gasser, L. Guillon, O. Cunrath, I. J. Schalk, Cellular organization of siderophore biosynthesis in Pseudomonas aeruginosa: Evidence for siderosomes. J. Inorg. Biochem. 148, 27–34 (2015).

55. V. Gasser, et al., In cellulo FRET-FLIM and single molecule tracking reveal the supra-molecular organization of the pyoverdine bio-synthetic enzymes in Pseudomonas aeruginosa. Q. Rev. Biophys. 53, e1 (2020).

56. C. Graf, M. Stankiewicz, G. Kramer, M. P. Mayer, Spatially and kinetically resolved changes in the conformational dynamics of the Hsp90 chaperone machine. EMBO J. 28, 602–613 (2009).

57. O. Genest, J. R. Hoskins, J. L. Camberg, S. M. Doyle, S. Wickner, Heat shock protein 90 from Escherichia coli collaborates with the DnaK chaperone system in client protein remodeling. Proc. Natl. Acad. Sci. U. S. A. 108, 8206–8211 (2011).

58. F. Tiburzi, F. Imperi, P. Visca, Intracellular levels and activity of PvdS, the major iron starvation sigma factor of Pseudomonas aeruginosa. Mol. Microbiol. 67, 213–227 (2008).

59. J. H. Seol, et al., The heat-shock protein HslVU from Escherichia coli is a protein-activated ATPase as well as an ATP-dependent proteinase. Eur. J. Biochem. 247, 1143–1150 (1997).

60. M. Bochtler, et al., The structures of HsIU and the ATP-dependent protease HsIU-HsIV. Nature 403, 800–805 (2000).

61. A.-R. Kwon, B. M. Kessler, H. S. Overkleeft, D. B. McKay, Structure and reactivity of an asymmetric complex between HslV and I-domain deleted HslU, a prokaryotic homolog of the eukaryotic proteasome. J. Mol. Biol. 330, 185–195 (2003).

62. A.-R. Kwon, C. B. Trame, D. B. McKay, Kinetics of protein substrate degradation by HslUV. J. Struct. Biol. 146, 141–147 (2004).

63. R. E. Burton, T. A. Baker, R. T. Sauer, Nucleotide-dependent substrate recognition by the AAA+ HslUV protease. Nat. Struct. Mol. Biol. 12, 245–251 (2005).

64. S. Tronnet, et al., Iron Homeostasis Regulates the Genotoxicity of Escherichia coli That Produces Colibactin. Infect. Immun. 84, 3358–3368 (2016).

65. G. L. Katumba, H. Tran, J. P. Henderson, The Yersinia High-Pathogenicity Island Encodes a Siderophore-Dependent Copper Response System in Uropathogenic Escherichia coli. mBio 13, e0239121 (2022).

66. L. E. Quadri, T. A. Keating, H. M. Patel, C. T. Walsh, Assembly of the Pseudomonas aeruginosa nonribosomal peptide siderophore pyochelin: In vitro reconstitution of aryl-4, 2-bisthiazoline synthetase activity from PchD, PchE, and PchF. Biochemistry 38, 14941–14954 (1999).

67. Y. Wang, D. Li, X. Huan, L. Zhang, H. Song, Crystallization and preliminary X-ray crystallographic analysis of a putative nonribosomal peptide synthase AmbB from Pseudomonas aeruginosa. Acta Crystallogr. Sect. F Struct. Biol. Commun. 70, 339–342 (2014).

68. N. Rojas Murcia, et al., The Pseudomonas aeruginosa antimetabolite L -2-amino-4-methoxy-trans-3-butenoic acid (AMB) is made from glutamate and two alanine residues via a thiotemplate-linked tripeptide precursor. Front. Microbiol. 6, 170 (2015).

69. S. Ernst, et al., Azetidomonamide and Diazetidomonapyridone Metabolites Control Biofilm Formation and Pigment Synthesis in Pseudomonas aeruginosa. J. Am. Chem. Soc. 144, 7676–7685 (2022).

70. B. Fauvet, et al., Bacterial Hsp90 Facilitates the Degradation of Aggregation-Prone Hsp70-Hsp40 Substrates. Front. Mol. Biosci. 8, 653073 (2021).

71. M. Corteggiani, et al., Uncoupling the Hsp90 and DnaK chaperone activities revealed the in vivo relevance of their collaboration in bacteria. Proc. Natl. Acad. Sci. U. S. A. 119, e2201779119 (2022).

72. T. Morán Luengo, M. P. Mayer, S. G. D. Rüdiger, The Hsp70-Hsp90 Chaperone Cascade in Protein Folding. Trends Cell Biol. 29, 164–177 (2019).

73. J. E. Cassat, E. P. Skaar, Iron in infection and immunity. Cell Host Microbe 13, 509–519 (2013).

74. Y. G. Santin, et al., In vivo TssA proximity labelling during type VI secretion biogenesis reveals TagA as a protein that stops and holds the sheath. Nat. Microbiol. 3, 1304–1313 (2018).

75. Y. Perez-Riverol, et al., The PRIDE database resources in 2022: a hub for mass spectrometry-based proteomics evidences. Nucleic Acids Res. 50, D543–D552 (2022).

