## Supporting Materials and Methods; Figures S1 to S5; Tables S1 to S4; SI References for "Bacterial Hsp90 promotes virulence factor production through maintenance of NRPS megaenzymes"

**This PDF file includes:**

Supporting Materials and Methods
Figures S1 to S5
Tables S1 to S4
SI References

**Other supporting materials for this manuscript include the following:**

Datasets S1 to S14

### Supporting Materials and Methods

#### Growth conditions

*E. coli* strains were grown in LB or DMEM media, and *P. aeruginosa* strains were grown in LB, iron-limiting CAA medium (Casamino acids vitamin assay, Gibco), or *Pseudomonas* isolation agar (PIA Difco laboratories-glycerol 0.5%) media (**Table S4**). *E. coli and P. aeruginosa* were cultured at the temperatures indicated in aerobic conditions with shaking at 180 rpm except if specified. When necessary, ampicillin (50 µg/mL), kanamycin (50 µg/mL), chloramphenicol (25 µg/mL) or streptomycin (50 µg/mL (*E. coli*), 2000 µg/mL (*P. aeruginosa*)), tetracycline (15 µg/mL (*E. coli*), 50 or 200 µg/mL (*P. aeruginosa*)) was added.

For proteomic analysis and quantitative RT-PCR, *E. coli* M1/5 WT or mutant strains were first grown overnight in LB medium at 37°C with shaking, then inoculated in DMEM. After 3h growth at 37°C with shaking, cultures were diluted to $OD_{600}$=0.03 in 10 mL DMEM and grown at 37°C in closed 14 mL tubes without shaking for 5 h.

For proteomic analysis, quantitative RT-PCR, extracellular pyoverdine measurement, and detection of Pvd proteins on SDS gels or western blot of *P. aeruginosa* grown under iron limiting conditions, cells were first grown overnight in LB medium and then overnight in CAA medium at 37°C. The following day, strains were diluted at $OD_{600}$=0.1 in CAA and cultured for 5 hours at the indicated temperature. For proteomic analysis in rich LB medium, *P. aeruginosa* strains were grown overnight at 37°C, inoculated at $OD_{600}$=0.1, and grown for 5h at 37°C.

#### Plasmid construction

All the plasmids and primers used in this study are listed in **Table S2** and **S3**, respectively. Cloning was performed either using restriction enzymes and T4 DNA ligase, sequence and ligation independent cloning (SLIC, (1)), or NEBuilder HiFi DNA Assembly kit. Cloned sequences were confirmed by DNA sequencing.

The following plasmids were used to construct *P. aeruginosa* strains by "sequence and ligation independent cloning" (SLIC) (1) except when indicated.

- The *P. aeruginosa* cis-complemented strain Δ$hsp90_{Pa}$::$hsp90_{Pa}$+ was generated by amplification of the $hsp90_{Pa}$ gene along with a 500 bp fragment corresponding to the putative promoter region of $hsp90_{Pa}$ and by cloning it into the pMiniCTX1 vector (2) yielding pMiniCTX1-$hsp90_{Pa}$. Mutation of the sequence coding for Hsp90$^{*E37A}$, was obtained by QuickChange site directed mutagenesis (Agilent).
- *P. aeruginosa* cis-complemented strains Δ$hslUV_{Pa}$::$hslUV_{Pa}$+ and Δ$hsp90_{Pa}$ Δ$hslUV_{Pa}$::$hslUV_{Pa}$ were generated by amplification of $hslUV_{Pa}$ genes along with a 500 bp fragment corresponding to the putative promoter region of $hslUV_{Pa}$ and by cloning it into the pMiniCTX1 vector (2) yielding pMiniCTX1-$hslUV_{Pa}$.
- To generate the *P. aeruginosa* $hsp90_{Pa}$ mutant strain, upstream and downstream 500 bp flanking regions of the $hsp90_{Pa}$ gene (PA1596) were cloned into the suicide pKNG101 vector (3) at the restriction sites *Bam*HI and *Spe*I, leading to the pKNG101-$hsp90_{Pa}$ plasmid. The same strategy was used to delete the $hslV_{Pa}$ (PA5053) and $hslU_{Pa}$ (PA5054) genes. The 500 bp upstream of $hslV_{Pa}$ sequence and the 500 bp downstream of *hslU* sequence were cloned into pKNG101, leading to the plasmid pKNG101-$hslUV_{Pa}$. These plasmids were transformed into the *E. coli* strain CC118λpir (4).
- To insert a 6His-tag sequence downstream of *pvdD* and *pvdJ* in *P. aeruginosa*, we cloned a sequence containing approximately 500 bp upstream of the stop codon of *pvdD* or *pvdJ*, the sequence coding for the 6His-tag, the stop codon, and approximately 500 bp downstream of *pvdD* or *pvdJ* in the pKNG101 plasmid, leading to the plasmids pKNG101-*pvdD*-6His and pKNG101- *pvdJ*-6His, respectively.
- To monitor translation of PvdI in *P. aeruginosa*, a sequence comprising the promoter of *pvdIJD* and the first 10 codons of *pvdI* were cloned in frame of the coding sequence of GFP (originated from pMiniCTX1-GFP) into pMiniCTX1 plasmid (2).

The following plasmids were used in *E. coli*. The pBAD33-, pBAD24-, pJF119EH- and pACYC184-based plasmids were constructed using NEBuilder HiFi DNA Assembly kit (NEB) using primers indicated in **Table S3**.

- To construct pACYC184-*clbB*::*lacZ,* a 686 bp sequence including the *clbR* transcriptional regulator, the putative promoter of *clbB*, RBS, ATG, and the first 45 nucleotides of *clbB* was cloned upstream of the *lacZ* gene (from amino acid 9) in pACYC184::*lacZ* at BamHI restriction site.
- To construct pBad24*hsp90*$_{Ec}$, the coding sequence of *hsp90*$_{Ec}$ was amplified by PCR, and cloned in the pBad24 vector at the *EcoRI* and *SalI* restriction sites.
- To construct pJF119EH -*hslUV*$_{Ec}$, the coding sequence of *hslUV*$_{Ec}$ was cloned in pJF119EH digested with SmaI and XbaI.
- To construct pBad33-*clbB*-6His, the coding sequence of *clbB* ended by a sequence coding for a 6-histidine tag were cloned in the pBad33 vector at the *Xba*I and *HindIII* restriction sites.
- To construct pBad33 *hsp90*$_{Pa}$, the coding sequence of *hsp90*$_{Pa}$ was amplified by PCR, and cloned in the pBad24 vector at the *SacI* and *XbaI* restriction sites

Plasmids were introduced by transformation in *E. coli* strains and by conjugation in *P. aeruginosa* using pRK2013 (5). The *E. coli* CC118λpir strain was used to propagate pKNG101 (3) derivatives, *E. coli* SM10 to propagate Mini-CTX1 (2) derivatives, while DH5α strains were used for other plasmids.

**Strain construction**

*E. coli* strains. The *hsp90*$_{Ec}$ *and hslV*$_{Ec}$ genes were deleted from the *E. coli* MC4100 strain by P1 transduction, using a lysate from *E. coli* JW0462 or *E. coli* JW3902 (Keio collection,(6)), yielding respectively the MC4100 Δ*hsp90*$_{Ec}$ strain or the MC4100 Δ*hslV*$_{Ec}$ strain.
The *hslV*$_{Ec}$ gene was deleted from the *E. coli* MC4100 Δ*hsp90*$_{Ec}$ strain by P1 transduction, using a lysate from *E. coli* JW3902 (Keio collection,(6)), yielding the MC4100 Δ*hsp90*$_{Ec}$Δ*hslV*$_{Ec}$ strain.

*P. aeruginosa* strains. Plasmids were transferred in *P. aeruginosa* strains by triparental conjugation using *E. coli* DH5α (for pMiniCTX1 derivatives) or the CC118λpir strain (for pKNG101 derivatives) as the donor and the conjugative properties of the helper plasmid pRK2013 as described before (7). The recombinant clones containing pMiniCTX1 derivatives inserted at the *attB* locus on the *P. aeruginosa* genome were selected on tetracycline-containing *Pseudomonas* isolation agar (PIA). In frame gene deletion mutants (PAO1 Δ*hsp90*$_{Pa}$, PAO1 Δ*hslUV*$_{Pa}$, PAO1 Δ*hslUV*$_{Pa}$Δ*hsp90*$_{Pa}$) and mutants producing PvdD-6His or PvdJ-6His were constructed by allelic exchange (8). Bacteria that have integrated the pKNG101_based plasmids into their genome were first selected on streptomycin-containing PIA. Then, isolation on Luria-Bertani (LB) plates containing 6% sucrose selected bacteria that have excised the plasmid, resulting in the deletion of the chromosomal target gene or in the insertion of the his tag sequence. Clones that became sucrose resistant and streptomycin sensitive were confirmed to contain the gene deletion or his tag insertion by PCR analysis.

**Proteomic analysis**

Pellets containing $6.10^7$ cells from each culture were collected after centrifugation. For *E. coli* samples and samples of *P. aeruginosa* grown in LB, cell pellets were resuspended in denaturing loading buffer, heat-denatured, and loaded on SDS-PAGE. Migration was stopped when the samples migrated as a thin band in the stacking part of the gel, then the bands were cut. Proteins contained in gel slices were reduced with DTT, alkylated with iodoacetamide and digested by Trypsin/LysC Mix from *P. aeruginosa* according to the protocol previously described (9), exception for *P. aeruginosa* samples treated by Trypsin Gold (Promega, Madison, USA). For samples of *P. aeruginosa* grown in CAA medium, the cells were lysed by sonication in 25 µL of lysis buffer (SDS % 50 mM Triethylammonium bicarbonate buffer, 0.5 mg/mL Dnase), and the supernatants were collected after centrifugation. The reduced and alkylated protein containing supernatants (performed as described here above for protein containing gel slices) were loaded on S-trap micro-columns (ProtiFi, New York, USA) according to the manufacturer's instructions to allow the trypsin digestion.

After preparation, extraction and drying in a speed vac, peptides were resuspended in a loading buffer (2% ACN, 0.1% TFA in H2O) and separated on a reversed phase column (EASY-Spray C18 PepMap 15 cm × 75 µm I.D, 100 Å pore size, 2 µm particle size, Thermo Fisher Scientific) at 40°C and 300 nL.min$^{-1}$ with a binary 90 min gradient. The gradient started with the mobile phases 98% A: 0.1% (v/v) formic acid in H2O and 2% B: 0.1% (v/v) formic acid in acetonitrile, increased to 25% B over the next 60 min, followed by a step gradient to 90% B for 6 min. Mass spectra were acquired on line in Data Dependent Acquisition mode (DDA) with a Q-orbitrap (Q-Exactive plus, Thermo Fisher Scientific). The Raw data were processed using either Proteome Discoverer 2.4 (Thermo Fisher Scientific) for protein identification or Maxquant 2.0.3.0 (10) for global quantitative proteomics searching with the databanks of the organism under study (for *E. coli* M1/5: chromosome CP053296 (4710 entries); plasmid CP053297 (38 entries); plasmid CP053298 (111 entries) (11); for *P. aeruginosa*: databank PAO1 strain downloaded from Uniprot on 26/12/2021 (7517 entries). The contaminant databank implemented in Maxquant was also searched. All parameters for LFQ quantitation were left as default and the match between run option was unabled. Resulting Proteingroups.txt files were exported to Perseus software platform (version 1.6.15.0) for statistical analysis (12). Briefly, after log 2 transformation of protein LFQ intensities, contaminant proteins, proteins matching in the reverse database, and proteins only identified by peptides that carry one or more modified amino acids were filtred out. Normal distribution of the data was then checked graphically. Proteins were considered as quantifiable if an intensity value was detected in above 50% of replicates in at least one condition (WT or mutants). Missing values were then imputed assuming a normal distribution of 0.3 width and 1.8 downshifts. Significant data points were determined using a t-test with a 250-permutation based FDR calculation (with 5% FDR and S0=0.1) and represented as a volcano plot. Differential proteins were considered significant if their fold change was >2 (Difference in $\log_2$ (FC) > 1 on the x axis) with $p<0.05$ ($-\log_{10}$(pvalue) > 1.3 on the y axis).

**RNA preparation and reverse transcription**

For *E. coli*, RNA was extracted and reverse transcribed into cDNA as described earlier (13). For *P. aeruginosa*, RNA was extracted with the "SV Total RNA Isolation System" kit from Promega before undergoing DNase treatment with the "Turbo DNA-free" kit (InvitrogenTM). The RNA quality was assessed by tape station 4200 system (Agilent). RNA was quantified spectrophotometrically at 260 nm (NanoDrop 1000; Thermo Fisher Scientific). For cDNA synthesis, 1 µg total RNA and 0.5 µg random primers (Promega) were used with the GoScript™ Reverse transcriptase (Promega) according to the manufacturer instruction.

**Quantitative Real-Time PCR**

Quantitative real-time PCR (qPCR) analyses were performed on a CFX96 Real-Time System (Bio-Rad). The reaction volume was 15 µL and the final concentration of each primer was 0.5 µM. The cycling parameters of the qPCR were 98°C for 2 min, followed by 45 cycles of 98°C for 5 s, 57°C for 10 s, 72°C for 1s. A final melting curve from 65°C to 95°C is added to determine the specificity of the amplification. To determine the amplification kinetics of each product, the fluorescence derived from the incorporation of EvaGreen into the double-stranded PCR products was measured at the end of each cycle using the SsoAdvanced™ Sybrgreen® 2X Kit (Bio-Rad, France). The results were analyzed using Bio-Rad CFX Maestro software, version 2.3 (Bio-Rad, France). For each point a technical duplicate was performed. The amplification efficiencies for each primer pairs were comprised between 80 and 100%. The *hcaT* and *cysG* genes were used as references for normalization for *E. coli* samples and the *16S, rpoD and rpsL genes* were used as references for normalization for *P. aeruginosa* samples. All primer pairs used for qPCR are reported in **Table S3**.

**Translational fusions**

*E. coli*. The pACYC184-ClbB::*lacZ*, pACYC184::*lacZ* and pACYC184 plasmids were introduced in MC4100, MC4100Δ*hsp90*$_{Ec}$, MC4100Δ*hslV*$_{Ec}$ and MC4100Δ*hsp90*$_{Ec}$Δ*hslV*$_{Ec}$ by electroporation. Cells were grown overnight in LB rich medium at 37°C. Strains were diluted 1/100 in LB and grown 2h30 at 37°C. β-galactosidase activity was measured using a modified Miller assay adapted for use in a Tecan Spark microplate reader as described previously (13).

*P. aeruginosa.* Pellets corresponding to 1ml of cells (approx. $OD_{600}$ 1.2) were obtained by centrifugation and resuspended in 200 µL PBS 1X. GFP fluorescence (Ex: 485 nm; Em: 535 nm) was measured with a microplate reader (Tecan), and the values were normalized to the corresponding $OD_{600}$. To remove endogenous fluorescence due to intracellular pyoverdine, the ratio fluorescence/$OD_{600}$ of WT or Δ$hsp90_{Ec}$ strain that do not produce GFP was subtracted to the ratio of WT or Δ$hsp90_{Ec}$ containing the mini-CTX1-GFP.

**Extracellular pyoverdine measurement**
Extracellular medium of 1 mL of cells ($OD_{600}$ around 1.2) were collected after two centrifugations at 13,000xg and transfer of 2/3 of the supernatant in a new tube after each centrifugation. Pyoverdine fluorescence (Ex: 400 nm; Em: 485 nm) was measured on 100 µL of the supernatant with a microplate reader (Tecan), and the values were normalized to the corresponding $OD_{600}$.

**Gene complementation:**
- For *hsp90* complementation, the WT and Δ$hsp90_{Ec}$ MG1655 *E. coli* strains containing pBAD33-*clbB*-6His (or pBAD33) and pBAD24-$hsp90_{Ec}$ (or pBAD24) were grown overnight in LB at 37°C, diluted 1/100 in LB, and incubated at 37°C. At $OD_{600}$=0.6, 0.2 % arabinose was added and cells were grown for 80 min. The same number of cells was collected, resuspended in loading buffer, heat-denatured, and proteins were separated on SDS-PAGE. The Coomassie InstantBlue® (abcam) was used for protein staining.
- For *hslV* complementation, the WT, Δ$hsp90_{Ec}$, and Δ$hsp90_{Ec}$Δ$hslV_{Ec}$ MG1655 *E. coli* strains containing pBAD33-*clbB*-6His and pJF119EH-$hslUV_{Ec}$ (or pJF119) were grown overnight in LB at 37°C, diluted 1/100 in LB supplemented with 100µM of IPTG, and incubated at 37°C. At $OD_{600}$=0.6, 0.2 % arabinose was added and cells were grown for 80 min. The same number of cells was collected, resuspended in loading buffer, heat-denatured, and proteins were separated on SDS-PAGE. The Coomassie InstantBlue® (abcam) was used for protein staining.

**ClbB detection by Western blot**
WT or mutant *E. coli* MG1655 strains containing the p*clbB*-6His plasmid were grown overnight in LB at 37°C, diluted 1/100 in LB, and incubated at 37°C. At $OD_{600}$=0.6, 0.02 % arabinose was added and cells were grown for 80 min. The same number of cells was collected, resuspended in loading buffer, heat-denatured, and proteins were separated on SDS-PAGE and transferred to nitrocellulose membranes. ClbB-6His was detected using a 6-His antibody (Invitrogen), and $Hsp90_{Ec}$ was detected with an anti-$Hsp90_{Ec}$ antibody (14). Band intensity was measured using ImageJ software and the amount of ClbB-6His measured in the WT strain was set to 100%.

**Detection of the Pvd proteins**
For whole cell extracts, an equal number of cells was centrifuged, and pellets were resuspended in denaturing Laemli buffer. Protein samples equivalent to 0.15 $OD_{600}$ units (anti-EF-Tu immunoblots) or 1.5 $OD_{600}$ units (anti-6His immunoblot) were loaded in each lane, separated on SDS-PAGE (8% and 8-20% polyacrylamide gel for Pvd proteins and EF-Tu respectively) and either stained with InstantBlue Coomassie Protein Stain (Abcam), or transferred to a nitrocellulose membrane. Proteins were revealed by Western blot and detected with monoclonal antibodies directed against 6His epitope-tag (Invitrogen, dilution 1:3300) and EF-Tu (Hycult®Biotech, dilution 1:10000). For anti 6His detection, manufacturer instructions were followed. Peroxidase-conjugated anti-Mouse IgGs (Sigma, dilution 1:5000) were used as secondary antibodies. The membranes were developed with homemade enhanced chemiluminescence assay and were scanned using ImageQuant TL analysis software (GE Healthcare Life sciences). Band intensity was measured using ImageQuant software and the amount measured in the WT strain was set to 100%.

**Detection of proteins involved in yersiniabactin production**
To visualize proteins involved in yersiniabactin production, WT and Δ*hsp90* M1/5 *E. coli* strains were grown overnight at 37°C in LB rich medium supplemented with 200 µM 2,2'-bipyridyl. Bacteria were then diluted 1/100 in LB medium supplemented with 200 µM of 2,2'-bipyridyl. As controls, 2,2'-bipyridyl was also omitted in all these steps. After overnight cultures, $5.10^7$ cells

from each culture were collected by centrifugation. The pellets were resuspended in loading buffer, heat-denaturated at 95°C for 10 min, and proteins were separated in SDS-PAGE that was stained with InstantBlue Coomassie Protein Stain (Abcam).

***P. aeruginosa* growth monitoring**

*P. aeruginosa* WT and $\Delta$*hsp90*$_{Pa}$ strains grown overnight in LB medium at 37°C were diluted to $OD_{600}$=1 in LB or CAA medium, and grown at 37°C or 42°C for 4 hours. These precultures were then used to inoculate 200 µL LB or CAA medium to $OD_{600}$=0.05 in 96-well plates. Growth was then monitored at 37°C or 42°C by measuring $OD_{600}$ with a microplate reader (Tecan).

### Supporting figures

Fig. S1

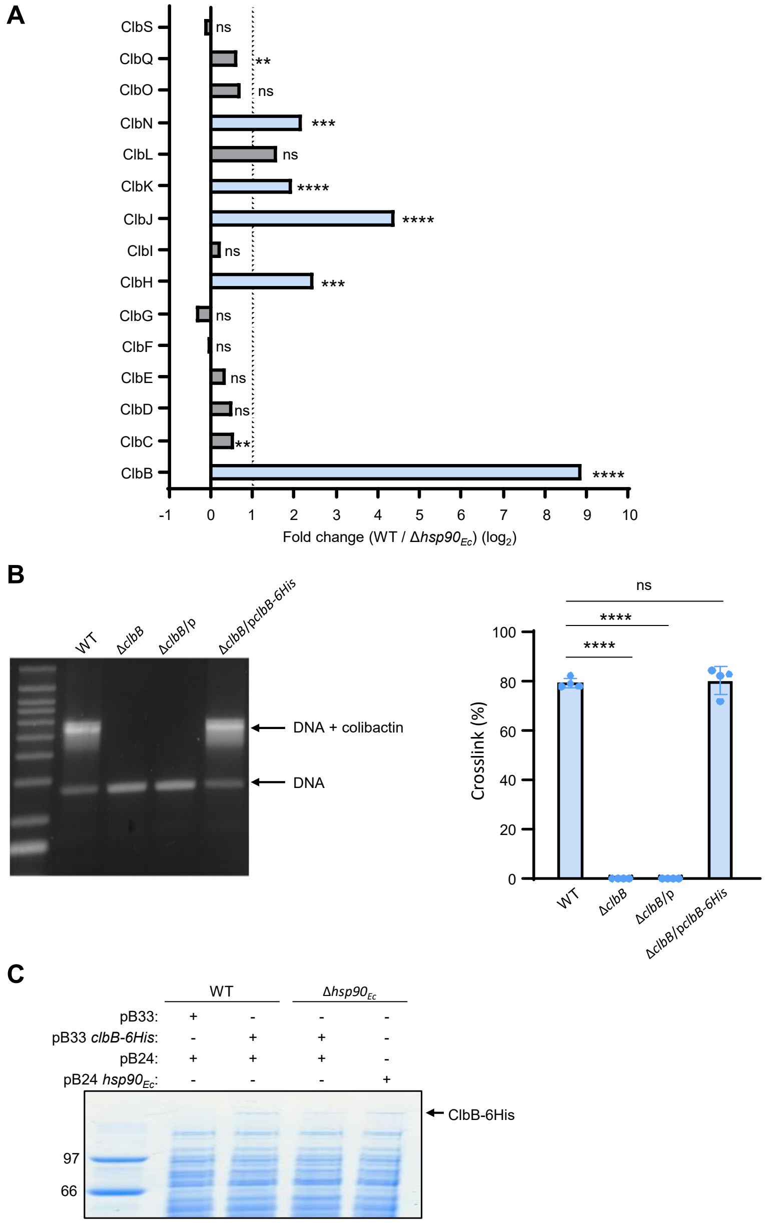

**Fig. S1. Hsp90$_{Ec}$ is involved in colibactin production. Controls related to Fig. 1.**

(**A**) Relative abundance of the proteins involved in colibactin biosynthesis that were detected in the mass spectrometry experiments in Fig. 1B. The ratio and their statistical analysis have been obtained from 4 biological replicates by the MaxQuant-Perseus software. The p-value associated with each ratio is indicated on the graph. P-values associated with each ratio indicate whether the differences are significant or not (****P ≤ 0.0001, ***P ≤ 0.001, **P ≤ 0.01, *P ≤ 0.05, ns: not significant).

(**B**) Functionality of ClbB with a 6His tag. ClbB with a 6-His tag was produced from a plasmid under the control of an arabinose promoter in WT, and Δ*clbB E. coli* Nissle strains. The functionality of the pBAD33-*clbB-6HIS* plasmid was confirmed by an assay in which the production of colibactin was examined through its DNA crosslinking activity. The assay was performed in four independent experiments, with the wild-type strain Nissle, Nissle Δ*clbB*::kana hosting the pBAD33 empty vector, or hosting the pBAD33-*clbB-6HIS* plasmid. The bacteria were inoculated at

$3.10^7$ bacteria/ml and cultivated in DMEM Hepes with 0.2% arabinose during 3.5 hours. Then 500 ng of linearized pUC19 plasmid DNA and 1mM EDTA were added, and the bacteria and DNA were further incubated during 40 minutes. The DNA was purified and analyzed by denaturing gel electrophoresis, as previously described (15). The DNA in the denatured and crosslinked DNA bands was quantified by image analysis using ImageJ. Results of one-way ANOVA tests indicate whether the differences measured are significant (****: p-value ≤ 0.0001) or not significant (ns, p-value > 0.05).
(**C**) Production of $Hsp90_{Ec}$ from a plasmid complements the reduced level of ClbB in the $\Delta$*hsp90*$_{Ec}$ strain. A pBAD24-*hsp90*$_{Ec}$ plasmid (or an empty pBAD24 vector, as negative control) was introduced in the *E. coli* MG1655 WT or $\Delta$*hsp90*$_{Ec}$ strain. The strains contained the pBAD33 *clbB*-6His plasmid (or a pBAD33 vector as control). Arabinose was added to induce *clbB* and *hsp90*$_{Ec}$ expression from the pBAD promoters. Same amount of proteins was separated on SDS-PAGE and visualized by Coomassie blue staining. ClbB migrates as the largest detectable protein in *E. coli*. This gel is representative of three independent experiments.

Fig. S2

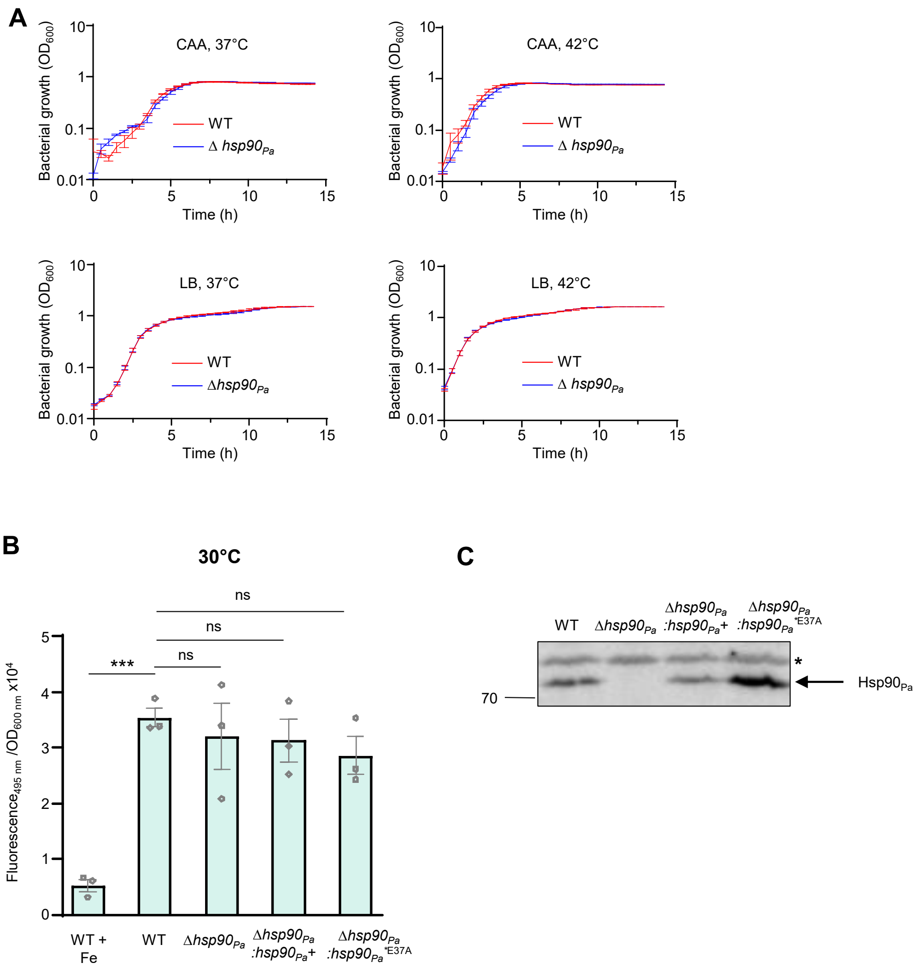

**Fig. S2. Hsp90 participates in pyoverdine production. Controls related to Fig. 2.**
(**A**) The absence of Hsp90 does not alter *P. aeruginosa* growth. *P. aeruginosa* WT, and Δ*hsp90* strains were grown in a microplate reader at 37°C and 42°C in LB or iron-limiting medium (CAA). Data from three replicates are shown as mean +/- SEM.
(**B**) Pyoverdine production measurement. *P. aeruginosa* WT, Δ*hsp90*, Δ*hsp90*:*hsp90*+, and Δ*hsp90*:*hsp90**$_{E37A}$ strains were grown at 30°C in iron-limiting medium (CAA) or, as a control in CAA supplemented with iron (WT + Fe). After 5 h of growth, pyoverdine was quantified by measuring fluorescence (excitation: 400 nm; emission: 485 nm) that was standardized to the number of cells ($OD_{600}$). Data from three biological replicates are shown as mean ± SEM. Results of one-way ANOVA tests indicate whether the differences measured are significant (***: p-value ≤ 0.001) or not significant (ns, p-value > 0.05).
(**C**) Hsp90 amount in *P. aeruginosa* strains used in the manuscript. *P. aeruginosa* WT, Δ*hsp90*:*hsp90*+, and Δ*hsp90*:*hsp90**$_{E37A}$ strains were grown at 37°C in iron-limiting medium (CAA). Hsp90 was detected with an $Hsp90_{Pa}$ antibody. The band marked with a star that is non-specifically revealed by the antibody was used as loading control. The Western blots shown are representative of 2 independent experiments.

Fig. S3

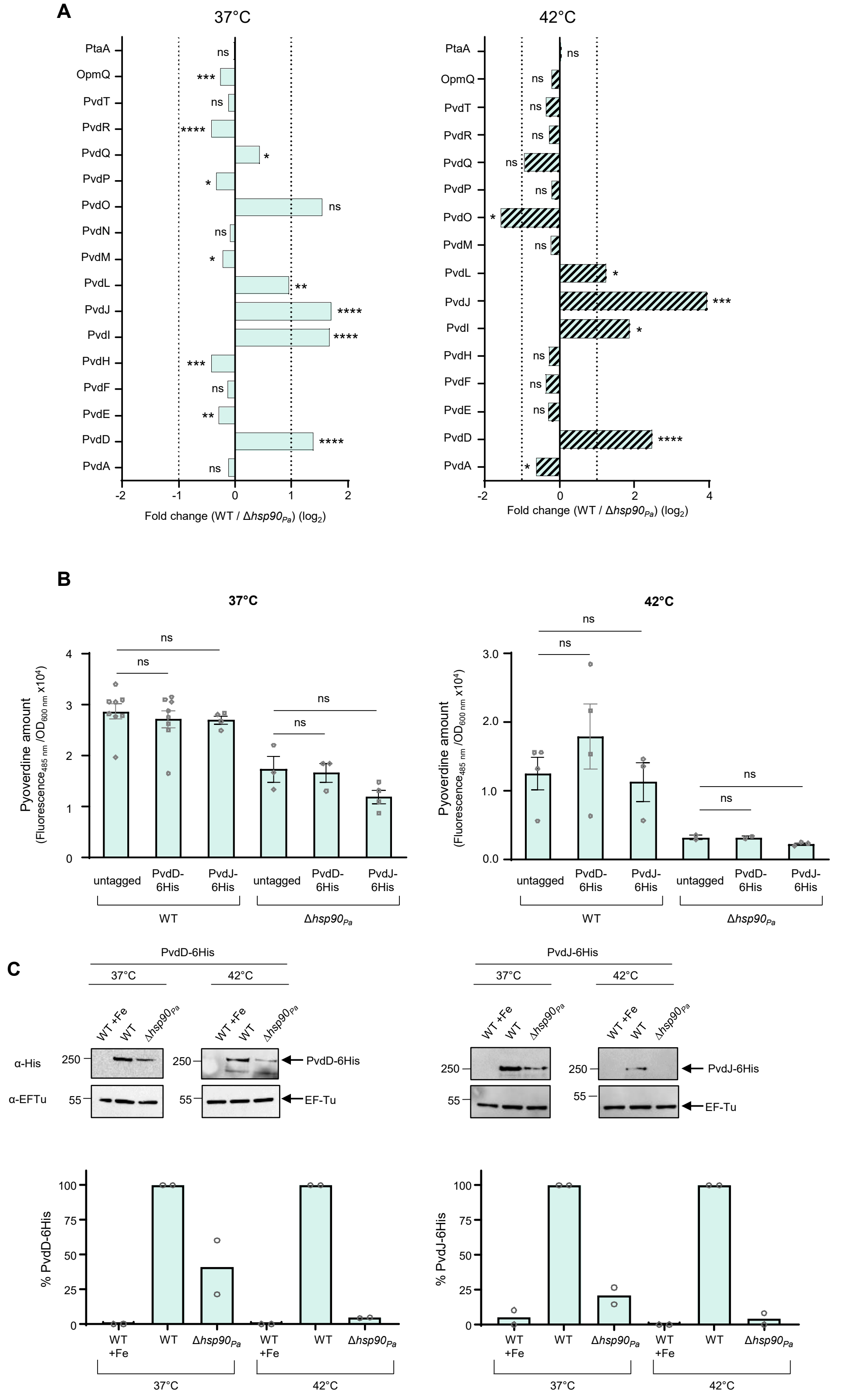
A
37°C
42°C
PtaA
OpmQ
PvdT
PvdR
PvdQ
PvdP
PvdO
PvdN
PvdM
PvdL
PvdJ
PvdI
PvdH
PvdF
PvdE
PvdD
PvdA
Fold change (WT / Δhsp90Pa) (log2)
B
37°C
42°C
Pyoverdine amount
(Fluorescence485 nm /OD600 nm x10⁴)
untagged
PvdD-6His
PvdJ-6His
WT
Δhsp90Pa
ns
C
PvdD-6His
PvdJ-6His
37°C
42°C
WT +Fe
WT
Δhsp90Pa
α-His
α-EFTu
PvdD-6His
PvdJ-6His
EF-Tu
% PvdD-6His
% PvdJ-6His
WT +Fe

**Fig. S3. Hsp90*Pa* stabilizes enzymes involved in pyoverdine production in *P. aeruginosa*. Controls related to Fig.3.**

(**A**) Relative abundance of the proteins involved in pyoverdine biosynthesis that were detected in the mass spectrometry experiments in Fig. 3A (37°C) and 3B (42°C). The ratio and their statistical analysis have been obtained from 4 biological replicates by the MaxQuant-Perseus software. The p-value associated with each ratio is indicated on the graph. P-values associated with each ratio indicate whether the differences are significant or not (****P ≤ 0.0001, ***P ≤ 0.001, **P ≤ 0.01, *P ≤ 0.05, ns: not significant).

(**B**) Addition of a 6His tag does not affect the functionality of PvdD and PvdJ. *P. aeruginosa* WT and Δ*hsp90* strains in which the *pvdD* and *pvdJ* genes were replaced by genes encoding 6His epitope-tagged proteins (PvdD-6His and PvdJ-6His) on the chromosome were grown at 37°C or 42°C in iron-limiting medium (CAA). After 5 h of growth, pyoverdine was quantified by measuring fluorescence (excitation: 400 nm; emission: 485 nm) that was standardized to the number of cells ($OD_{600}$). Data from three biological replicates are shown as mean ± SEM. Results of one-way ANOVA tests indicate that the differences measured are not significant (ns, p-value > 0.05).

(**C**) Replicate and quantification of the immunoblots shown in Figure 3E. Quantification of the amount of PvdD-6His (left) and PvdJ-6His (right) was performed from 2 independent Western blots using ImageJ software. The amount of PvdD-6His or PvdJ-6His measured in the wild-type strain was set to 100%.

Fig. S4

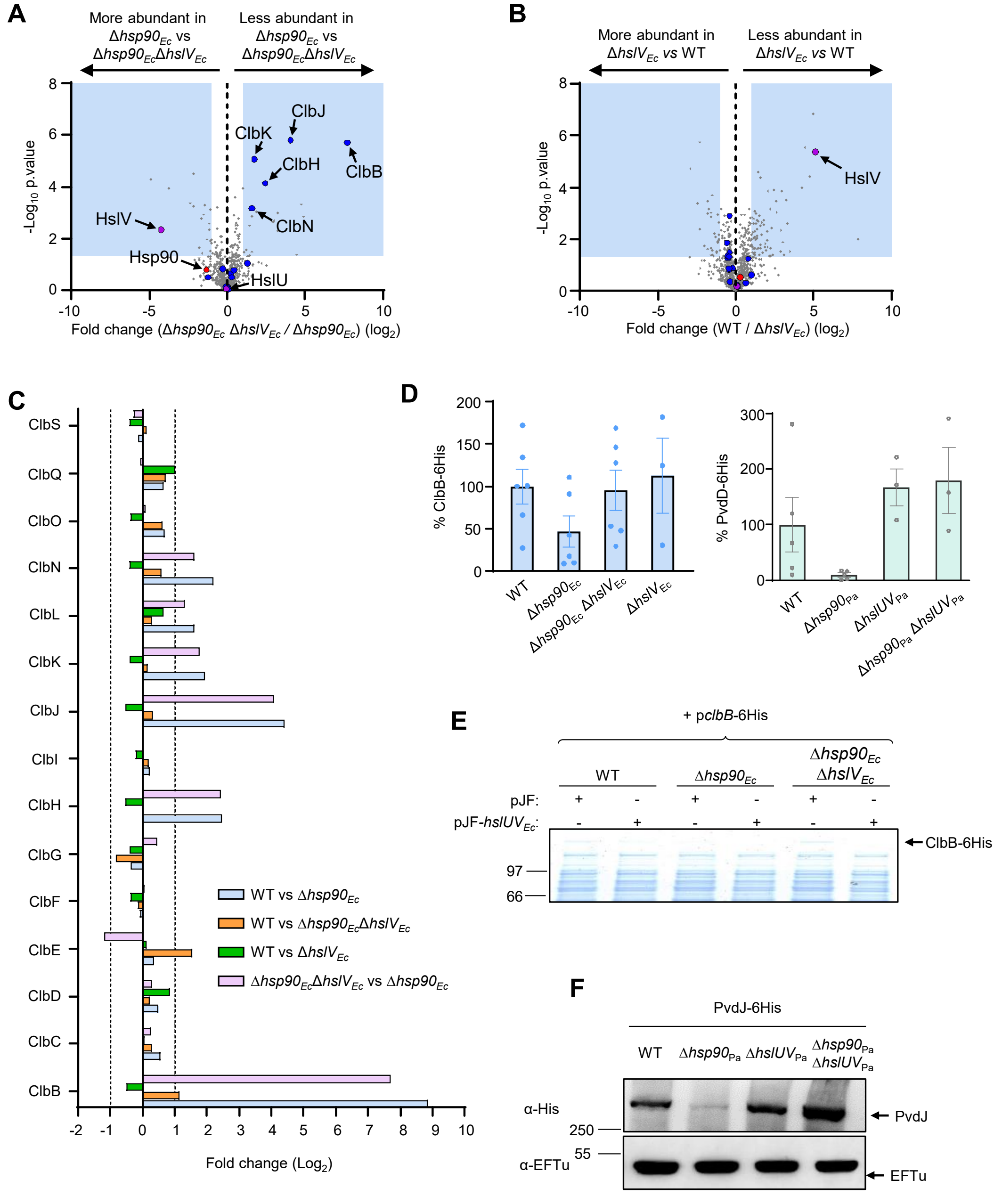

A
More abundant in Δhsp90Ec vs Δhsp90EcΔhslVEc
Less abundant in Δhsp90Ec vs Δhsp90EcΔhslVEc
-Log10 p.value
ClbJ
ClbK
ClbH
ClbB
HslV
ClbN
Hsp90
HslU
Fold change (Δhsp90Ec ΔhslVEc / Δhsp90Ec) (log2)
B
More abundant in ΔhslVEc vs WT
Less abundant in ΔhslVEc vs WT
-Log10 p.value
HslV
Fold change (WT / ΔhslVEc) (log2)
C
ClbS
ClbQ
ClbO
ClbN
ClbL
ClbK
ClbJ
ClbI
ClbH
ClbG
ClbF
ClbE
ClbD
ClbC
ClbB
WT vs Δhsp90Ec
WT vs Δhsp90EcΔhslVEc
WT vs ΔhslVEc
Δhsp90EcΔhslVEc vs Δhsp90Ec
Fold change (Log2)
D
% ClbB-6His
WT
Δhsp90Ec
Δhsp90Ec ΔhslVEc
ΔhslVEc
% PvdD-6His
WT
Δhsp90Pa
ΔhslUVPa
Δhsp90Pa ΔhslUVPa
E
+ pclbB-6His
WT
Δhsp90Ec
Δhsp90Ec ΔhslVEc
pJF:
pJF-hslUVEc:
ClbB-6His
97
66
F
PvdJ-6His
WT
Δhsp90Pa
ΔhslUVPa
Δhsp90Pa ΔhslUVPa
α-His
250
PvdJ
α-EFTu
55
EFTu

Fig. S4 (continued)

G

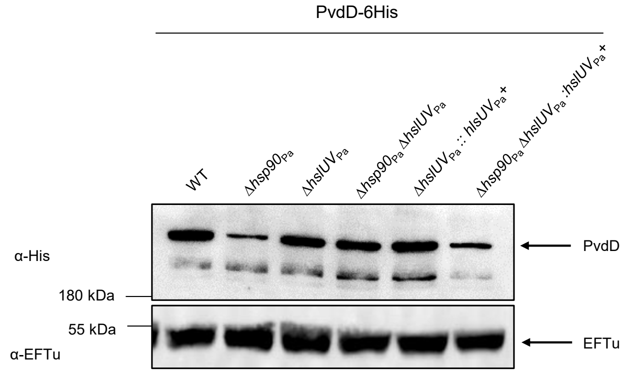

H

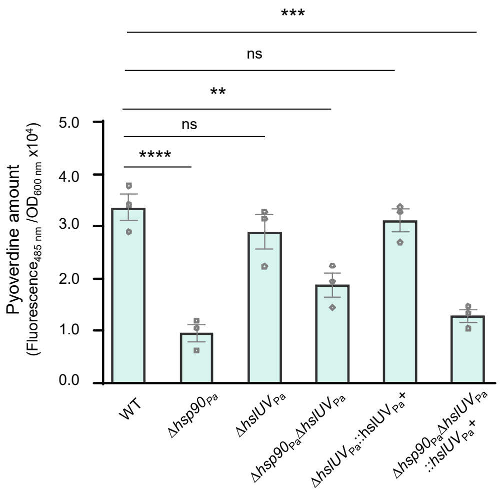

**Fig. S4. Interplay between Hsp90 and HslUV to control the level of NRPS. Controls related to Fig.4.**

(**A, B**) Global comparative quantitative proteomics analysis of *E. coli* $\Delta hsp90_{Ec}$ vs $\Delta hsp90_{Ec}\Delta hslV_{Ec}$ (A), and $\Delta$*hslV* vs WT (B) strains. The protein content of the two strains grown at 37°C was analyzed by LFQ mass spectrometry, and the relative abundance of proteins is reported. The X axis indicates the protein fold change ($log_2$) between the two strains and the Y axis indicates the p-value ($-log_{10}$) of this difference. Regions of the graph framed in light blue indicate proteins whose abundance varied significantly between the two strains (fold change >2; p-value <0.05). Proteins of interest are pointed by arrows. Hsp90$_{Ec}$ is colored in red, and proteins involved in colibactin biosynthesis in blue. The experiments were performed with 4 biological replicates for each strain, and the results were analyzed with MaxQuant/Perseus software suite.

(**C**) Relative abundance of proteins involved in colibactin biosynthesis measured in Fig. 1B, 4A, and S4A-B. p-values associated with each ratio indicate whether the differences are significant or not (****P ≤ 0.0001, ***P ≤ 0.001, **P ≤ 0.01, *P ≤ 0.05, ns: not significant).

(**D**) Quantification of the Western blots shown in Fig. 4C. The intensity of the bands corresponding to ClbB (left) and PvdD (right) was quantified using ImageQuant. The amount of ClbB or PvdD measured in the WT strain was set to 100%. Data from three or more biological replicates are shown as mean ± SEM.

(**E**) Production of HslUV$_{Ec}$ from a plasmid leads to reduced levels of ClbB. The pJF-$hslUV_{Ec}$ was introduced in *E. coli* MG1655 WT, $\Delta hsp90_{Ec}$ and $\Delta hsp90_{Ec}\Delta hslV_{Ec}$ containing a plasmid allowing ClbB production. Expression of $hslUV_{Ec}$ was induced by addition of IPTG, followed by induction of *clbB* by arabinose. Same amount of proteins were separated by SDS-PAGE and visualized by Coomassie blue staining. ClbB migrates as the largest detectable protein in *E. coli*. This gel is representative of three independent experiments.

(**F**) Absence of the $HslUV_{Pa}$ protease increases the level of PvdJ in the Δ*hsp90* strain. *P. aeruginosa* WT, $\Delta hsp90_{Pa}$, $\Delta hslUV_{Pa}$, and $\Delta hsp90_{Pa}\Delta hslUV_{Pa}$ strains carrying a chromosomal encoded C-terminal 6-His epitope tagged PvdJ (PvdJ-6His, 241 kDa) were grown at 42°C in iron-deficient medium (CAA). PvdJ was detected by immunoblot with a 6His antibody. Antibody against EF-Tu (43.3 kDa) was used as loading control. Molecular weight markers (kDa) are indicated on the left. The Western blots shown are representative of two independent experiments

(**G**) Complementation of the $\Delta hsp90_{Pa}\,\Delta hslUV_{Pa}$ strain by production of $HslUV_{Pa}$. The $hslUV_{Pa}$ genes were reintroduced on the chromosome of *P. aeruginosa* leading to strains $\Delta hslUV_{Pa}::hlsUV_{Pa}$+ and $\Delta hsp90_{Pa}$ $\Delta hslUV_{Pa}::hslUV_{Pa}$+. Strains as indicated and carrying a chromosomal encoded C-terminal 6-His epitope tagged PvdD (PvdD-6His, 274.5 kDa) were grown at 42°C in iron-deficient medium (CAA). PvdD was detected by immunoblot with a 6His antibody. Antibody against EF-Tu (43.3 kDa) was used as loading control. Molecular weight markers (kDa) are indicated on the left. The Western blots shown are representative of three independent experiments.

(**H**) Pyoverdine amount quantification in strains as in G. Strains were grown at 42°C in iron-limiting medium (CAA). After 5 h of growth, pyoverdine was quantified by measuring fluorescence (excitation: 400 nm; emission: 485 nm) that was standardized to the number of cells ($OD_{600}$). Data from three biological replicates are shown as mean ± SEM. Results of one-way ANOVA tests indicate whether the differences measured are significant (****: p-value ≤ 0.0001; ***: p-value ≤ 0.001; **: p-value ≤ 0.01) or not significant (ns, p-value > 0.05).

Fig. S5

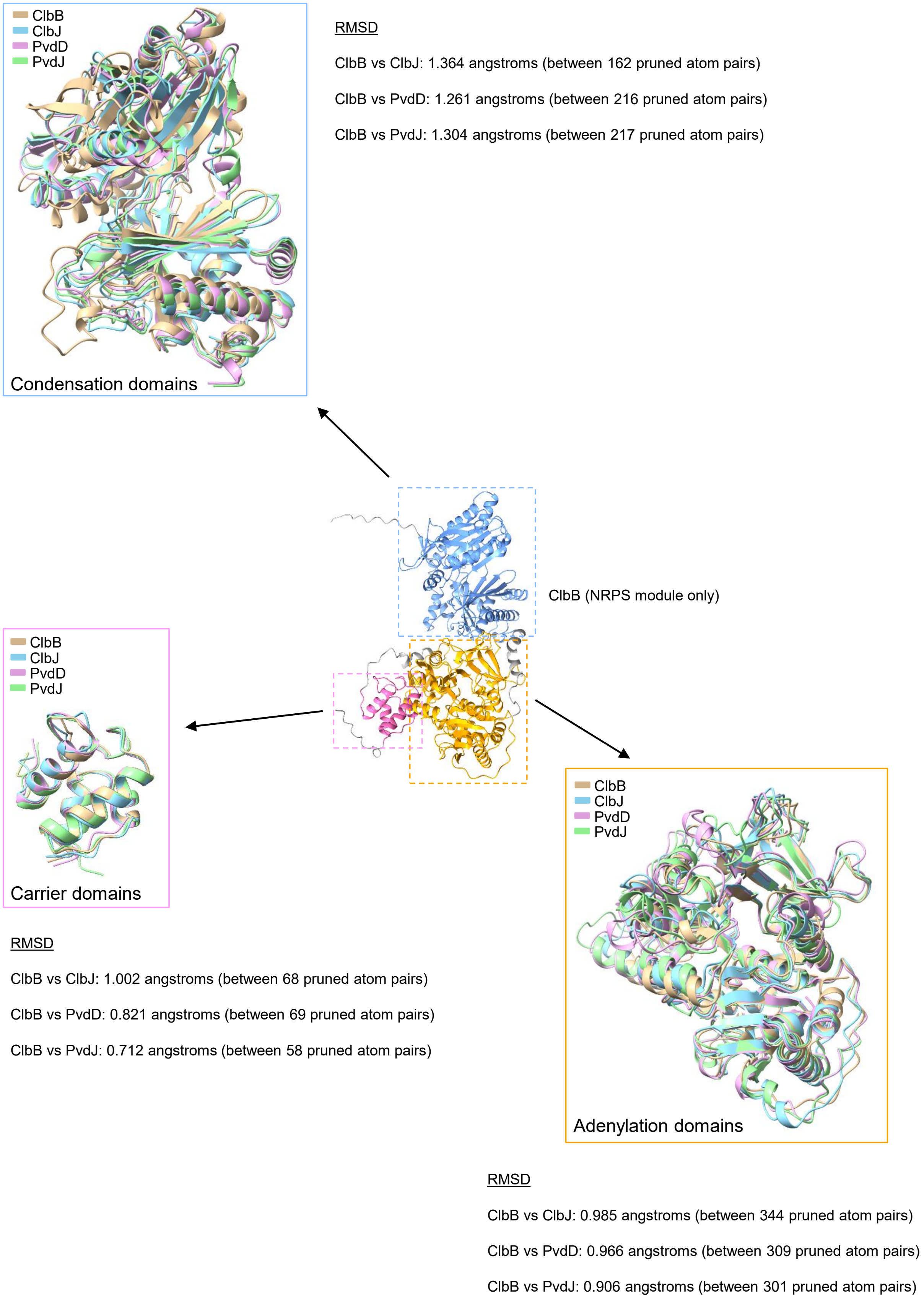

ClbB
ClbJ
PvdD
PvdJ
Condensation domains
RMSD
ClbB vs ClbJ: 1.364 angstroms (between 162 pruned atom pairs)
ClbB vs PvdD: 1.261 angstroms (between 216 pruned atom pairs)
ClbB vs PvdJ: 1.304 angstroms (between 217 pruned atom pairs)
ClbB (NRPS module only)
ClbB
ClbJ
PvdD
PvdJ
Carrier domains
RMSD
ClbB vs ClbJ: 1.002 angstroms (between 68 pruned atom pairs)
ClbB vs PvdD: 0.821 angstroms (between 69 pruned atom pairs)
ClbB vs PvdJ: 0.712 angstroms (between 58 pruned atom pairs)
ClbB
ClbJ
PvdD
PvdJ
Adenylation domains
RMSD
ClbB vs ClbJ: 0.985 angstroms (between 344 pruned atom pairs)
ClbB vs PvdD: 0.966 angstroms (between 309 pruned atom pairs)
ClbB vs PvdJ: 0.906 angstroms (between 301 pruned atom pairs)

**Fig. S5. Structure prediction of the NRPS module of some megasynthases.**
Structure predictions by Alphafold3. In the center of the figure, Alphafold3 structure prediction of the NRPS module of ClbB (residues 1 to 1083). The condensation domain (17-473) is blue, the adenylation domain (498-970) is orange, and the carrier domain (994-1069) is pink. The regions connecting these domains are in gray.
In boxes: overlay of the condensation domains, adenylation domains, and carrier domains from the first NRPS module of ClbB, ClbJ, PvdD, and PvdJ. Condensation domains: ClbB: 17-479; ClbJ: 43-477; PvdD: 46-486; PvdJ: 16-459. Adenylation domains: ClbB: 498-970; ClbJ: 508-976; PvdD: 508-999; PvdJ: 482-969. Carrier domains: ClbB: 994-1069; ClbJ: 998-1073; PvdD: 1019-1094; PvdJ: 989-1064. The domain limits were obtained from InterPro website (https://www.ebi.ac.uk/interpro/). The images and RMSD calculations were prepared using the ChimeraX software (version 1.5).

**Table S1: Strains used in this study**

| Strain | Genotype / Description | Source or reference |
|---|---|---|
| ***Escherichia coli*** | | |
| **DH5α** | *F$^-$ φ80dlacZ*ΔM15 Δ*(lacZYA-argF)U169 recA1 endA1 hsdR17($r_k^-$ $m_k^+$) phoA supE44 thi-1 gyrA96 relA1* | Laboratory Stock |
| **CC118λpir** | Δ*(ara-leu) ara*D Δ*lacX74 galE galK phoA20 thi-1 rspE rpoB argE (Am) recA1 λpir* | (4) |
| **1047/pRK2013** | Mobilizing strain for conjugation, *KanR, oriColE1 RK2- mob+ tra+* | (16) |
| **MG1655** | Wild-type *E. coli* strain; F- λ- ilvG- rfb-50 rph-1 | (17) |
| **M1/5** | Wild-type *E. coli* strain; colibactin genotoxin producer. | (11) |
| **M1/5 Δ*hsp90*$_{Ec}$** | Deletion of the *hsp90*$_{Ec}$ gene in strain M1/5 | (18) |
| **M1/5 Δ*hslV*$_{Ec}$** | Deletion of the *hslV*$_{Ec}$ gene in strain M1/5 | (18) |
| **M1/5 Δ*hsp90*$_{Ec}$Δ*hslV*$_{Ec}$** | Deletion of the *hsp90*$_{Ec}$ and *hslV*$_{Ec}$ genes in strain M1/5 | (18) |
| **MC4100** | Wild-type *E. coli* strain; F$^-$ *araD139* Δ*(argF-lac) U169 rpsL150 relA1 flb-5301 fruA25 deoC1 ptsF25* | (19) |
| **MC4100 Δ*hsp90*$_{Ec}$** | Deletion of the *hsp90*$_{Ec}$ gene in strain MC4100 | This study |
| **MC4100 Δ*hslV*$_{Ec}$** | Deletion of the *hslV*$_{Ec}$ gene in strain MC4100 | This study |
| **MC4100 Δ*hsp90*$_{Ec}$Δ*hslV*$_{Ec}$** | Deletion of the *hsp90*$_{Ec}$ and *hslV*$_{Ec}$ genes in strain MC4100 | This study |
| **MG1655** | Wild-type *E. coli* strain; F- λ- ilvG- rfb-50 rph-1 | (17) |
| **MG1655 Δ*hsp90*$_{Ec}$** | Deletion of the *hsp90*$_{Ec}$ gene in strain MG1655 | (14) |
| **MG1655 Δ*hslV*$_{Ec}$** | Deletion of the *hslV*$_{Ec}$ gene in strain MG1655 | This study |
| **MG1655 Δ*hsp90*$_{Ec}$Δ*hslV*$_{Ec}$** | Deletion of the *hsp90*$_{Ec}$ and *hslV*$_{Ec}$ genes in strain MG1655 | This study |
| **JW0462** | Deletion of the *hsp90*$_{Ec}$ gene, Keio collection | (6) |
| **JW3902** | Deletion of the *hslV*$_{Ec}$ gene, Keio collection | (6) |
| **Nissle 1917** | Wild-type E. coli strain (Mutaflor, DSM 6601, serotype O6:K5:H1) | (20) |
| **Nissle Δ*clbB*** | Deletion of the *clbB* gene in strain Nissle 1917 | (20) |
| ***Pseudomonas aeruginosa*** | | |
| **PAO1 WT** | Wild-type | Laboratory Stock |
| **PAO1 Δ*hsp90*$_{Pa}$** | *hsp90*$_{Pa}$ deletion mutant | This study |
| **PAO1 Δ*hslUV*$_{Pa}$** | *hslUV*$_{Pa}$ deletion mutant | |
| **PAO1 Δ*hslUV*$_{Pa}$Δ*hsp90*$_{Pa}$** | *hslUV*$_{Pa}$ *and hsp90*$_{Pa}$ deletion mutant | |
| **PAO1 Δ*hsp90*$_{Pa}$::*hsp90*$_{Pa}$ +** | Deletion mutant of the *hsp90*$_{Pa}$ gene complemented with *hsp90*$_{Pa}$ gene at the *att*B site on the chromosome, Tc$^R$. | |

| | |
|---|---|
| **PAO1 Δ$hsp90_{Pa}$::$hsp90_{Pa*}^{E37A}$** | Deletion mutant of the $hsp90_{Pa}$ gene complemented with $hsp90_{Pa*}^{E37A}$ gene at the *att*B site on the chromosome, $Tc^R$. |
| **PAO1 Δ$hslUV_{Pa}$::$hslUV_{Pa}$** | Deletion mutant of $hslUV_{Pa}$ genes complemented with $hslUV_{Pa}$ genes at the *att*B site on the chromosome, $Tc^R$. |
| **PAO1 Δ$hsp90_{Pa}$ Δ$hslUV_{Pa}$::$hslUV_{Pa}$** | $hslUV_{Pa}$ *and* $hsp90_{Pa}$ deletion mutant complemented with $hslUV_{Pa}$ genes at the *att*B site on the chromosome, $Tc^R$. |
| **PAO1 WT PvdD-6His** | Insertion of a 6-His tag sequence downstream of the *pvdD* gene at the locus. |
| **PAO1 Δ$hsp90_{Pa}$ PvdD-6His** | |
| **PAO1 Δ$hslUV_{Pa}$ PvdD-6His** | |
| **PAO1 Δ$hsp90_{Pa}$Δ$hslUV_{Pa}$ PvdD-6His** | |
| **PAO1 WT PvdJ-6His** | Insertion of a 6-His tag sequence downstream of the *pvdJ* gene at the locus. |
| **PAO1 Δ$hsp90_{Pa}$ PvdJ-6His** | |
| **PAO1 Δ$hslUV_{Pa}$ PvdJ-6His** | |
| **PAO1 Δ$hsp90_{Pa}$Δ$hslUV_{Pa}$ PvdJ His** | |
| **PAO1 WT miniCTX1-GFP** | Wild-type strain with the *gfp* gene at the *attB* site on the chromosome, $Tc^R$ . |
| **PAO1 Δ$hsp90_{Pa}$ miniCTX1-GFP** | Deletion mutant of the $hsp90_{Pa}$ gene with *gfp* gene at the *att*B site on the chromosome, $Tc^R$. |
| **PAO1 Δ$hslUV_{Pa}$ miniCTX1-GFP** | Deletion mutant of $hslUV_{Pa}$ genes with *gfp* gene at the *att*B site on the chromosome, $Tc^R$. |
| **PAO1 Δ$hsp90_{Pa}$ Δ$hslUV_{Pa}$ miniCTX1-GFP** | $hslUV_{Pa}$ *and* $hsp90_{Pa}$ deletion mutant with *gfp* gene at the *att*B site on the chromosome, $Tc^R$. |
| **PAO1 WT miniCTX1-*pvdI*-GFP** | Wild-type strain with the *gfp* gene under the control of *pvdI* transcriptional and translation signals at the *attB* site on the chromosome, $Tc^R$ . |
| **PAO1 Δ$hsp90_{Pa}$ miniCTX1-*pvdI*-GFP** | Deletion mutant of $hsp90_{Pa}$ with the *gfp* gene under the control of *pvdI* transcriptional and translation signals at the *attB* site of the chromosome, $Tc^R$ . |
| **PAO1 Δ$hslUV_{Pa}$ miniCTX1-GFP** | Deletion mutant of $hslUV_{Pa}$ genes with the *gfp* gene under the control of *pvdI* transcriptional and translation signals at the *attB* site of the chromosome, $Tc^R$ . |
| **PAO1 Δ$hsp90_{Pa}$ Δ$hslUV_{Pa}$ miniCTX1-GFP** | $hslUV_{Pa}$ *and* $hsp90_{Pa}$ deletion mutant with the *gfp* gene under the control of *pvdI* transcriptional and translation signals at the *attB* site of the chromosome, $Tc^R$ . |

**Table S2: Plasmids used in this study**

| Plasmid | Description | Origine |
|---|---|---|
| **pBAD33** | pBAD33 plasmid. Arabinose inducible. CmR. | (21) |
| **pBAD33-*clbB-6HIS*** | pBAD33 plasmid carrying *clbB* gene with a 6-His tag sequence at the C-term extremity | This study |
| **pACYC184** | pACYC184 plasmid. CmR. | (22) |
| **pACYC184 :: *lacZ*** | *lacZ* gene cloned in the pACYC184 plasmid; no promoter upstream of *lacZ*. | (23) |
| **pACYC184-pClbB :: *lacZ*** | Transcriptional and translational signals of *clbB* cloned upstream and in frame with *lacZ*. | This study |
| **pBAD33-*hsp90*$_{Ec}$** | pBAD33 plasmid carrying *hsp90*$_{Ec}$ gene. | (24) |
| **pBAD33-*hsp90*$_{Pa}$** | pBAD33 plasmid carrying *hsp90*$_{Pa}$ gene. | This study |
| **pBAD24** | pBAD24 plasmid. Arabinose inducible. AmpR. | (21) |
| **pJF119EH** | Vector containing the Ptac (ApR) promoter with the pBR origin of replication | (25) |
| **pJF119EH-*hslUV*$_{Ec}$** | pJF119EH plasmid carrying *hslU*$_{Ec}$ *and hslV*$_{Ec}$ gene. | This study |
| **pMiniCTX1** | Non-replicative vector containing the *attP* site for integration into the *attB* site on the *P. aeruginosa* chromosome, Ω-FRT-*attP*-MCS, *ori*, *int*, and *oriT*, TetR | (2) |
| **pMiniCTX1-*hsp90*$_{Pa}$** | MiniCTX1 vector containing the *hsp90*$_{Pa}$ gene under the control of *its own* promoter | Laboratory Stock |
| **pMiniCTX1-*hsp90*$_{Pa}$$^{*E37A}$** | MiniCTX1 vector containing the *hsp90*$_{Pa}$ $^{*E37A}$ gene under the control of *hsp90*$_{Pa}$ promoter | This study |
| **pMiniCTX1-*hslUV*$_{Pa}$** | MiniCTX1 vector containing *hslUV*$_{Pa}$ genes under the control of *hslUV*$_{Pa}$ promoter | This study |
| **pRK2013** | *tra*$^{+}$, *mob*$^{+}$, *ori*$_{colE1}$, Kan$^{R}$ | (16) |
| **pKNG101** | *oriR6K, mobRK2, sacBR+, SmR (suicide vector).* | (16) |
| **pKNG101-*hsp90*$_{Pa}$** | pKNG101 vector containing 500 base pairs upstream and downstream of the *hsp90*$_{Pa}$ gene from *P. aeruginosa.* | This study |
| **pKNG101-*hslUV*$_{Pa}$** | pKNG101 vector containing 500 base pairs upstream and downstream of the *hslUV*$_{Pa}$ gene from *P. aeruginosa.* | |
| **pKNG101-*pvdD*-His** | pKNG101 vector containing 500 base pairs upstream (containing the 6His tag) and downstream of the 3' extremity of *pvdD*. | |
| **pKNG101- *pvdJ*-His** | pKNG101 vector containing 500 base pairs upstream (containing the 6His tag) and downstream of of the 3' extremity of *pvdJ*. | |
| **pMiniCTX1-GFP** | MiniCTX1 vector containing the *gfp* gene without promoter. | (26) |
| **pMiniCTX1 *pvdI*-GFP** | MiniCTX1 vector containing transcriptional and translational signals of *pvdI* cloned upstream and in frame with *gfp*. | This study |

**Table S3: Oligonucleotides used in this study**

| Oligonucleotide | Sequence | Purpose |
|---|---|---|
| **OG1332** | CGACCACACCCGTCCTGTTTAGATAATCTCATTCCTGT | To generate pACYC184-pClbB :: *lacZ* |
| **OG1333** | GTAAAACGACACGCGTGTCCATCTTATTAC | |
| **OG1334** | GGACACGCGTGTCGTTTTACAACGTCGTGA | |
| **EF086** | ACGATGCGTCCGGCGTAGATTATTATTATTTTTGACACCAG | |
| **OG733** | GAA GGATCC CCCTGGTGGCCGGCCAGGCTCAGG | To generate pKNG101-*hsp90*$_{Pa}$ |
| **OG734** | CGGGGCTACTGC CGCTCCAAGCTCCATCAATGACA | |
| **OG735** | GAGCTTGGAGCG GCAGTAGCCCCGACAAAGCCCGCT | |
| **OG736** | GAA ACTAGT ACACGAAGGCCTTCAGCTCGGCGC | |
| **OG946** | GGGCTAGCGAATTCGAGCTCGGAAGGAGATATACATATGGAT AATACCTCTGGAGATTTTCC | To generate pBad33-*clbB*-6His |
| **OG947** | GCCAGATAGCATGCAGCGCATGTTCCATGTCACTATGCAC | |
| **OG948** | GTGCATAGTGACATGGAACATGCGCTGCATGCTATCTGGC | |
| **OG949** | GGTGCTGTGCATACACCAGCGAGCAACACGATAAAGTCCC | |
| **OG950** | GGGACTTTATCGTGTTGCTCGCTGGTGTATGCACAGCACC | |
| **OG951** | GGTCGACTCTAGAGGATCCCCGGTTAGTGGTGGTGGTGGTG GTGATGCAAAGACGTGTGACTGCGCGCAG | |
| **OG1170** | TGGGCTAGCGAATTCGAGCTCAAATAAGGAAAATTTCATGGTG GCAGAATCCCGCACCGG | To generate pBad33-*hsp90*$_{Pa}$ |
| **OG1171** | TTGCATGCCTGCAGGTCGACTCTAGATTATTGCTCCAGCAGG CGCTTC | |
| **OG1174** | TAGCTGATTCACACGACGAA | qPCR (*clbB*) |
| **OG1175** | GTCTTCCACAATGTCCTTGC | |
| **OG1176** | GCGTCAACAGGAATACATCG | qPCR (*clbH*) |
| **OG1177** | AAAGCGGCAAACAGAATACC | |
| **OG1180** | TTGGATATGCAGCGTACCTT | qPCR (*clbJ*) |
| **OG1181** | GGCAATCAGCGGAATCATAC | |
| **OG1182** | CTGTGCTTGGGAGATGTTTG | qPCR (*clbK*) |
| **OG1183** | GCTTGTCTGGGTTGTTCATC | |
| **OG1184** | AGTCTCTCATGGCAATCTGG | qPCR (*clbN*) |
| **OG1185** | GCGTAGTGATGGTCAAATCG | |
| **OG1190** | GCTGGCACTGCTGACA | qPCR (*hcaT*) |
| **OG1191** | CGCCGAGCCAATGACA | |
| **OG1211** | GTCGCATCTTCTGTAACGTG | qPCR (*cysG*) |
| **OG1212** | GCAGCAGTGATTCGAGTTTT | |
| **OG1209** | CAGAATTCGAGCTCGGTAACTCTGTATTCGTAACCAAGG | To generate pJF119EH-*hslUV*$_{Ec}$ |
| **OG1210** | GCATGCCTGCAGGTCGACTTATAGGATAAAACGGCTCAG | |
| ***hsp90*$_{Ec}$ Up** | TAGAATTCATGAAAGGACAAGAAACTCGT | To generate pBad 24-*hsp90*$_{Ec}$ |
| ***hsp90*$_{Ec}$ Down** | TAGTCGACTCAGGAAACCAGCAGCTGGTTC | |
| ***hslUV*$_{Pa}$ up F** | CAGGTCGACGGATCCCCGGGGCCAAGCCGAAGTACGAGTT | To generate pKNG-*hslUV*$_{Pa}$ |
| ***hslUV*$_{Pa}$ up R** | CGGTGGTCACAAGGGGGAAATCTCCACAC | |
| ***hslUV*$_{Pa}$ Down F** | TCCCCCTTGTGACCACCGCCGCCGCCACG | |
| ***hslUV*$_{Pa}$ Down R** | TATGCATCCGCGGGCCCGGGGGCGAATGTTGCAAATGACG | |
| ***hslUV*$_{Pa}$ UU** | TTCCTGATGAAGCTCGAACC | |
| ***hslUV*$_{Pa}$ DD** | CGGCTACTGCTCGACTACCA | |
| ***Aco2*** | GCTTGATATCGAATTCTTAAGCGGACAGCTCCACCA | To generate miniCTX1-*hsp90*$_{Pa}$ |
| ***Aco3*** | CGGGCTGCAGGAATTCATACCTTGCACCTGGACCTG | |
| ***hslUV CTX For*** | CCCCCGGGCTGCAGGAATTCCGTGCGCGCGCAGATCATCC | |

| | | |
|---|---|---|
| ***hslUV CTX Rev*** | TGTCCCGCTACATCCTTTGAGAATTCGATATCAAGCTTAT | To generate miniCTX1-$hslUV_{Pa}$ |
| ***pvdD his up F*** | CAGGTCGACGGATCCCCGGGGTCAGGGTCGAAGCGTTCT | To generate pKNG-*pvdD*-His |
| ***pvdD his up R*** | GTGATGGTGATGGTGATGGCGCCCGGCACGCTCCAGGG | |
| ***pvdD his down F*** | CATCACCATCACCATCACTGATCTCAGGTCCGTTCCGT | |
| ***pvdD down R*** | TATGCATCCGCGGGCCCGGGGAAATCTCGGGTGAACTGGC | |
| ***pvdD UU*** | CAACCTGGCGATGGATGTC | |
| ***pvdJ his up F*** | CAGGTCGACGGATCCCCGGGCAGGTGAAAATCCGAGGCTT | To generate pKNG-*pvdJ*-His |
| ***pvdJ his up R*** | GTGATGGTGATGGTGATGGGAAATCAGTTTTTCAAGTTCATCG G | |
| ***pvdJ his down F*** | CATCACCATCACCATCACTAAGAGGCGGTAGCGTGCAA | |
| ***pvdJ down R*** | TATGCATCCGCGGGCCCGGGCCTCGACTATAGGCGGCATA | |
| ***pvdJ UU*** | TTTCGTGCCGGATCCCTTT | |
| ***PpvdIup(Bam)*** | AGCTCGGTACCCGGGGATCCGAGGCTAACGGTAGGTTAGG | To generate miniCTX1-*pvdI*-GFP |
| ***PpvdI-ATGdown*** | AAGTTCTTCTCCTTTACTGGCGAGCTTGAGGGAATCTT | |
| ***pvdI-ATG-GFPup*** | GATTCCCTCAAGCTCGCCAGTAAAGGAGAAGAACTTTT | |
| ***CTXGFPdown (Spe1)*** | GTGGCGGCCGCTCTAGAAACTAGTGGATCGTACGAATG | |
| $hsp90_{Pa}$ ***E37A F** | ATCTTCCTCCGCGCGCTGATTTCCAAC | QuickChange |
| $hsp90_{Pa}$ ***E37A R** | GTTGGAAATCAGCGCGCGGAGGAAGAT | |
| ***pvdD* RT Up** | GAAGGCATTGGCTGTCCTGC | qPCR *P. aeruginosa* |
| ***pvdD* RT Down** | CACTGTCGCTCCTGGGCAAT | |
| ***pvdJ* RT Up** | TTCAATCCCAGGCCGACTC | |
| ***pvdJ* RT Down** | AACGATGCCCAGATACCGTC | |
| ***pvdI* RT Up** | TACGAGTTCAGCATCGAGCC | |
| ***pvdI* RT Down** | GAGGATATTGAAGCCGCCGA | |
| ***pvdL* RT Up** | TCCAGTACACCTCCGGTTCA | |
| ***pvdL* RT Down** | CCGGACGTTCGAAGTAG | |
| ***pvdS* RT Up** | GGAACAACTGTCTACCGCCA | |
| ***pvdS* RT Down** | TGAAGAACGCATCCTGGACC | |
| $hsp90_{Pa}$ **RT Up** | TCTACAGCGCCTTCATCGTC | |
| $hsp90_{Pa}$ **RT Down** | CTCCTCGCCCTTTTTCAGGT | |
| ***16S* RT Up** | CAGCTCGTGTCGTGAGATGT | |
| ***16S* RT Down** | GATCCGGACTACGATCGGTT | |
| ***rpoD RT For*** | GCGCAACAGCAATCTCGTCT | |
| ***rpoD RT Rev*** | ATCCGGGGCTGTCTCGAATA | |
| ***rpsL RT For*** | GCAAGCGCATGGTCGACAAGA | |
| ***rpsL RT Rev*** | CGCTGTGCTCTTGCAGGTTGTGA | |

**Table S4: Media used**

| Medium | Description |
|---|---|
| **LB (Luria-Bertani)** | Rich medium : Bacto-tryptone (10 g/L), yeast extract (5 g/L), NaCl (5 g/L) |
| **DMEM** | Dulbecco's Modified Eagle Medium, Gibco |
| **M9** | Minimum medium : M9 1X ($KH_2PO_4$, 3 g/L, NaCl 0.5 g/L, $Na_2HPO_4$ 6.78 g/L, $NH_4Cl$ 1 g/L), 0,5%, casamino acids, $MgSO_4$ 1 mM, $CaCl_2$ 0,2 mM, 0,2% Glucose. |
| **CAA (Casamino acids)** | Iron-limiting medium : commercial powder Difco$^{TM}$ « Casamino acids, Vitamin Assay », $MgSO_4$ 2 mM, $K_2HPO_4$ 8,4 mM. |
| **PIA** | Selective medium : commercial powder Difco$^{TM}$, 0,5% glycerol |

### SI References

1. J.-Y. Jeong, *et al.*, One-step sequence- and ligation-independent cloning as a rapid and versatile cloning method for functional genomics studies. *Appl. Environ. Microbiol.* **78**, 5440–5443 (2012).
2. T. T. Hoang, A. J. Kutchma, A. Becher, H. P. Schweizer, Integration-proficient plasmids for Pseudomonas aeruginosa: site-specific integration and use for engineering of reporter and expression strains. *Plasmid* **43**, 59–72 (2000).
3. K. Kaniga, I. Delor, G. R. Cornelis, A wide-host-range suicide vector for improving reverse genetics in gram-negative bacteria: inactivation of the blaA gene of Yersinia enterocolitica. *Gene* **109**, 137–141 (1991).
4. M. Herrero, V. de Lorenzo, K. N. Timmis, Transposon vectors containing non-antibiotic resistance selection markers for cloning and stable chromosomal insertion of foreign genes in gram-negative bacteria. *J. Bacteriol.* **172**, 6557–6567 (1990).
5. D. H. Figurski, D. R. Helinski, Replication of an origin-containing derivative of plasmid RK2 dependent on a plasmid function provided in trans. *Proc. Natl. Acad. Sci. U. S. A.* **76**, 1648–1652 (1979).
6. T. Baba, *et al.*, Construction of Escherichia coli K-12 in-frame, single-gene knockout mutants: the Keio collection. *Mol. Syst. Biol.* **2**, 2006.0008 (2006).
7. T. G. Sana, A. Laubier, S. Bleves, Gene transfer: conjugation. *Methods Mol. Biol. Clifton NJ* **1149**, 17–22 (2014).
8. H. P. Schweizer, T. T. Hoang, An improved system for gene replacement and xylE fusion analysis in Pseudomonas aeruginosa. *Gene* **158**, 15–22 (1995).
9. Y. G. Santin, *et al.*, In vivo TssA proximity labelling during type VI secretion biogenesis reveals TagA as a protein that stops and holds the sheath. *Nat. Microbiol.* **3**, 1304–1313 (2018).
10. S. Tyanova, T. Temu, J. Cox, The MaxQuant computational platform for mass spectrometry-based shotgun proteomics. *Nat. Protoc.* **11**, 2301–2319 (2016).
11. A. Wallenstein, *et al.*, ClbR Is the Key Transcriptional Activator of Colibactin Gene Expression in Escherichia coli. *mSphere* **5**, e00591-20 (2020).
12. S. Tyanova, J. Cox, Perseus: A Bioinformatics Platform for Integrative Analysis of Proteomics Data in Cancer Research. *Methods Mol. Biol. Clifton NJ* **1711**, 133–148 (2018).
13. N. J. Maillot, F. A. Honoré, Byrne, Deborah, V. Méjean, O. Genest, Cold adaptation in the environmental bacterium Shewanella oneidensis is controlled by a J-domain co-chaperone protein network. *Commun. Biol.* **2**, 323 (2019).
14. O. Genest, *et al.*, Uncovering a region of heat shock protein 90 important for client binding in E. coli and chaperone function in yeast. *Mol. Cell* **49**, 464–473 (2013).
15. N. Bossuet-Greif, *et al.*, The Colibactin Genotoxin Generates DNA Interstrand Cross-Links in Infected Cells. *mBio* **9**, e02393-17 (2018).
16. D. H. Figurski, D. R. Helinski, Replication of an origin-containing derivative of plasmid RK2 dependent on a plasmid function provided in trans. *Proc. Natl. Acad. Sci. U. S. A.* **76**, 1648–1652 (1979).
17. F. R. Blattner, *et al.*, The complete genome sequence of Escherichia coli K-12. *Science* **277**, 1453–1462 (1997).
18. C. Garcie, *et al.*, The Bacterial Stress-Responsive Hsp90 Chaperone (HtpG) Is Required for the Production of the Genotoxin Colibactin and the Siderophore Yersiniabactin in Escherichia coli. *J. Infect. Dis.* **214**, 916–924 (2016).
19. M. J. Casadaban, Transposition and fusion of the lac genes to selected promoters in Escherichia coli using bacteriophage lambda and Mu. *J. Mol. Biol.* **104**, 541–555 (1976).
20. T. Pérez-Berezo, *et al.*, Identification of an analgesic lipopeptide produced by the probiotic Escherichia coli strain Nissle 1917. *Nat. Commun.* **8**, 1314 (2017).
21. L. M. Guzman, D. Belin, M. J. Carson, J. Beckwith, Tight regulation, modulation, and high-level expression by vectors containing the arabinose PBAD promoter. *J. Bacteriol.* **177**, 4121–4130 (1995).

22. A. C. Chang, S. N. Cohen, Construction and characterization of amplifiable multicopy DNA cloning vehicles derived from the P15A cryptic miniplasmid. *J. Bacteriol.* **134**, 1141–1156 (1978).
23. H. Baaziz, *et al.*, ChrASO, the chromate efflux pump of Shewanella oneidensis, improves chromate survival and reduction. *PloS One* **12**, e0188516 (2017).
24. M. Corteggiani, *et al.*, Uncoupling the Hsp90 and DnaK chaperone activities revealed the in vivo relevance of their collaboration in bacteria. *Proc. Natl. Acad. Sci. U. S. A.* **119**, e2201779119 (2022).
25. J. P. Fürste, *et al.*, Molecular cloning of the plasmid RP4 primase region in a multi-host-range tacP expression vector. *Gene* **48**, 119–131 (1986).
26. A. Becher, H. P. Schweizer, Integration-proficient Pseudomonas aeruginosa vectors for isolation of single-copy chromosomal lacZ and lux gene fusions. *BioTechniques* **29**, 948–950, 952 (2000).
